# Mutations in disordered proteins as early indicators of nucleic acid changes triggering speciation

**DOI:** 10.1101/867648

**Authors:** Sergio Forcelloni, Andrea Giansanti

**Author notes:** **Corresponding Author:** Sergio Forcelloni, Sapienza University of Rome, Department of Physics, P.le A. Moro 5, 00185 Roma, Italy. **E-mail:**.

## Abstract

Intrinsically disordered proteins are characterized by unusual sequence composition, structural flexibility, and functional spectra. These properties play an essential role in fostering protein evolution and in the formation of complex cellular pathways, especially in multicellular organisms. In this study, we analyze the role of different structural variants of proteins in speciation processes. Firstly, we separate human and mouse proteomes (taken as a reference) in three variants of disorder: ordered proteins (ORDPs), structured proteins with intrinsically disordered protein regions (IDPRs), and intrinsically disordered proteins (IDPs). Secondly, we compare the DNA divergence with the corresponding protein divergence, by confronting human and mouse coding sequences (separated in ORDPs, IDPRs, and IDPs) with their homologs from 26 eukaryotes. As a general rule, we find that IDPs phenotypically diverge earlier than ORDPs and IDPRs. ORDPs diverge later but are phenotypically more reactive to nucleotide mutations than IDPRs and IDPs. We suggest that *IDPs may be involved in the early stages of the speciation process*, likely connected to their functional spectra, mainly related to nucleic acid binding and transcription factors. In contrast, *ORDPs may be essential in accelerating further phenotypic divergence.*

## 1. Introduction

It is quite known that under the chromosomal hypothesis the accumulation, in the genomes of prospective parents, of mutations that make their offspring sterile (because meiosis cannot proceed), might trigger speciation [1]. In this work, we address the question whether the genes coding for intrinsically disordered proteins selectively act, in the genomes of a group of mammalian species, as speciation genes. Intrinsically disordered proteins (IDPs) and proteins with long intrinsically disordered regions (IDPRs) are abundantly present in all proteomes, including viruses [2]. Interestingly, the content of IDPs/IDRs tends to increase with organism complexity [3]; approximately 25 – 30% of eukaryotic proteins are predicted to be mostly disordered and more than half of eukaryotic proteins have long disordered regions [4].

Comprehensive analyses have shown that proteins with loose packing, low degree of tertiary interactions, low compactness, and disorder are prone to have high evolvability (i.e., the ability of proteins to adopt new functions within the same fold or changing the fold) [5,6]. This is clearly reflected in the unusual properties of IDPs/IDPRs. Firstly, mutations in disordered regions cause smaller stability changes than those in ordered regions [7]. Secondly, the specific order of amino acids in the sequence of disordered proteins is less conserved than in well-structured ones, thus showing higher rates of accepted point mutations, insertions, and deletions in their sequence as long as they maintain enough disorder-prone amino-acids [8]. Therefore, the weaker impact of mutations in intrinsically disordered proteins and regions may have advantages for protein evolvability, in the gain of new functions.

During species divergence, populations accumulate genetic differences in both coding and noncoding regions, which ultimately can lead to genetic conflicts, and hence to hybrid sterility or inviability [1,9]. Several computational and experimental lines of evidence have provided two general evolutionary insights about genes which are responsible for speciation, commonly referred to as ‘speciation genes’ [10]. First, they are fast-evolving, likely driven by positive selection prompted by genetic conflicts, rather than by adaptation to environmental conditions. Second, many of the identified interspecific incompatibilities are caused by genes encoding DNA-binding proteins and transcription factors. In this context, recent studies have shown that both these features characterize intrinsically disordered proteins [11,12], thus suggesting a central role of these proteins in the speciation process.

Although all the above observations point to IDPs as highly evolvable proteins and to their genes as prospective speciation genes it is not well explored the question whether proteins with different percentages of disordered residues differently contribute to the speciation process. To address this question, we rely on the recent operational classification of protein disorder by Deiana et al. [12]. Thus, we distinguish three broad protein classes, characterized by different structural, functional, and evolutionary properties [12,13]. Namely: i) ordered proteins (ORDPs); ii) structured proteins with intrinsically disordered protein regions (IDPRs); iii) intrinsically disordered proteins (IDPs). In a nutshell, IDPs enrich only a few classes, functions, and processes: *nucleic acid-binding proteins, chromatin-binding proteins, transcription factors*, and *developmental processes.* In contrast, IDPRs are spread over several functional protein classes and GO annotations, partly shared with ORDPs. Moreover, we observed that ORDPs and IDPs proteins are more subject to natural selection than IDPRs, whereas IDPRs are more subject to mutational bias [13]. We anticipate that, from the data shown in this work, ORDPs and IDPs, though subject to high selective evolutionary pressure, are characterized by different mutational rates; higher in IDPs.

Based on these findings and motivated by Wolfe and Sharp [14], we compared the DNA divergences with the corresponding protein divergences, in several mammalian species, by considering separately ORDPs, IDPRs, and IDPs. Notably, differential trends among these protein variants were observed in the rate of nucleotide substitutions and phenotypic divergence, paving the way for a general interpretation of their roles in the speciation processes.

## 2. Results

### 2.1. ORDPs, IDPRs, and IDPs are characterized by different rates of evolution

Human genes were separated following the variant of disorder of the corresponding proteins (i.e., ORDPs, IDPRs, and IDPs) (see Materials and Methods). Each human gene was confronted with the homologous gene in Mus Musculus. For each pair of genes, we compared the base substitutions (DNA divergence) with the amino acid substitutions (protein divergence). At variance with nucleotide substitutions, amino acid substitutions are expected to result in phenotypic differences. This reflects the degeneracy of the genetic code so that many mutations in coding sequence are silent, i.e., they do not change the amino acid sequence encoded. Finally, we repeated this protocol for each gene in each variant of disorder, thus characterizing the rate of phenotypic divergence of each variant of disorder as a function of the nucleotide divergence. In Fig. 1 each point corresponds to an individual gene and, since DNA divergence increases with time, the X-axis can be conceived as a time axis [15].

**Figure 1.**
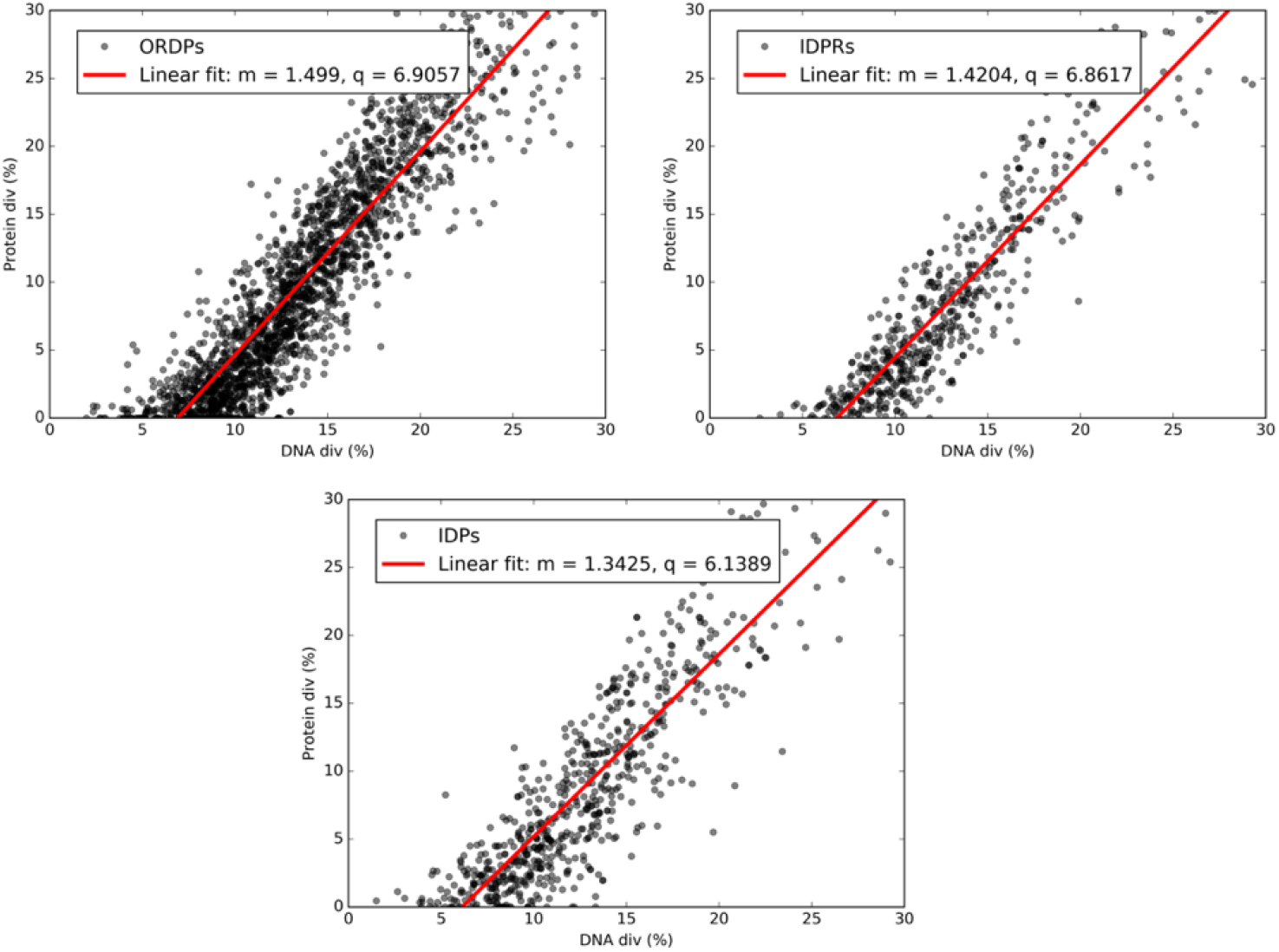
Example of DNA divergence vs. Protein divergence plot. Relationship between nucleotide (DNA div) and amino acid (Protein div) sequence divergence among genes compared between human and mouse. Each point corresponds to an individual gene. In each panel, we report the best-fit line in red, together with the associated values of the slope (m) and the intercept (q) in the legend.

As expected, the extents of protein and DNA sequence divergence are linearly correlated. Therefore, if a gene displays a low DNA divergence, then the corresponding protein will have zero or a few amino acid substitutions. Conversely, if a gene displays a high DNA divergence, then the encoded protein will have many amino acid substitutions.

In genes encoding highly conserved proteins, DNA divergence exceeds protein divergence, because silent synonymous substitutions are permitted in these genes. On the other hand, in genes encoding for less conserved proteins, protein divergence often exceeds DNA divergence. This fact is evident considering the following example: a codon with a single point mutation at position 2 (causing an amino acid change) has 33.3% of nucleotide divergence but 100% amino acid divergence.

Interestingly, all the intercepts in Fig. 1 are significantly different from 0. Thus, it appears that some proteins have remained unchanged, although the corresponding DNA sequences have diverged a little (genes on the X-axis). Presumably, any organisms with non-synonymous mutations have been negatively selected and, therefore, no mutations are found in modern organisms. Conversely, organisms with synonymous mutations that do not change the encoded protein sequence have not been counter-selected so severely [15].

To test the differences between the linear regressions associated with the variants of disorder, we used the protocol described in Materials and Methods. All the linear regressions in Fig. 1 are significantly different and, therefore, characterized by different intercepts and slopes.

Interpretations of X-axis intercept values would range from zero, when there is an equal probability that divergence was initiated at the protein and nucleic acid level, to increasing positive values reflecting increasing expectations that nucleic acid level differences triggered *initial* divergence. Our results provide evidence that after the primary nucleic acid level difference has triggered the speciation process, specific variants of disorder will contribute before others. Notably, the values of intercepts of ORDPs (6.9) and IDPs (6.14) suggest that the latter is triggered earlier by the nucleic acid changes than the former. In other words, ORDPs are more resistant to nucleic acid changes that will eventually recruit them to play a role in furthering the speciation process.

As regards to slopes, these values provide an estimation of the rate of phenotypic divergence as a function of nucleotide divergence. Thus, the steeper slope for ORDPs (1.54) than for IDPs (1.27) suggests that *once recruited*, nucleotide changes in the former play a stronger phenotypic role than nucleotide changes in the latter. At the same time, the less steep slope for IDP-encoding genes shows that they are freer to accept nucleic acid changes and especially synonymous mutations, which are typically neutral and reflect broad mutational processes acting also on non-genic genomic regions. In this context, it is worth noting that this ability of IDPs to evolve neutrally – a measure of robustness to mutations – has been related to high designability and facilitates evolutionary innovation on large time-scales [6,16,17].

### 2.2. IDPs trigger the speciation process, whereas ORDPs accelerate the phenotypic divergence

To further confirm these considerations, we generalize this analysis by confronting progressively human coding sequences (separated in ORDPs, IDPRs, and IDPs) with their homologs from 26 eukaryotes. All cases are similar to Fig. 1, with a well-defined linear relationship associated with each variant of disorder (see Figs S1). Thus, for each species in comparison with human and each variant of disorder, we retrieve the intercept (*q_variant_*) and the slope (*m_variant_*) of the best-linear fit. To confront the values of intercepts and slopes associated with the variants, we decided to rescale them as follows:

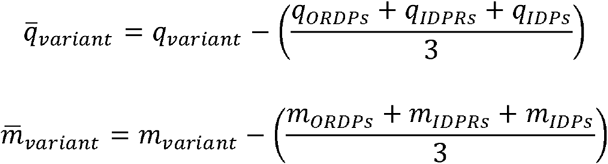

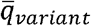 and 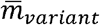 are the rescaled values of intercept and slope, with respect to the arithmetic mean of intercepts and slopes of all variants of disorder (amounts in brackets). In this framework, a 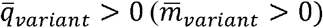 means that the intercept (slope) of the variant is *higher* than the average intercept (slope) of all variants; conversely, a 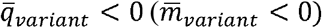 means that the intercept (slope) of the variant is less than the average intercept (slope) of all variants. In Figs. 2 and 3, we show the normalized values of intercepts and slopes obtained from all comparisons.

**Figure 2.**
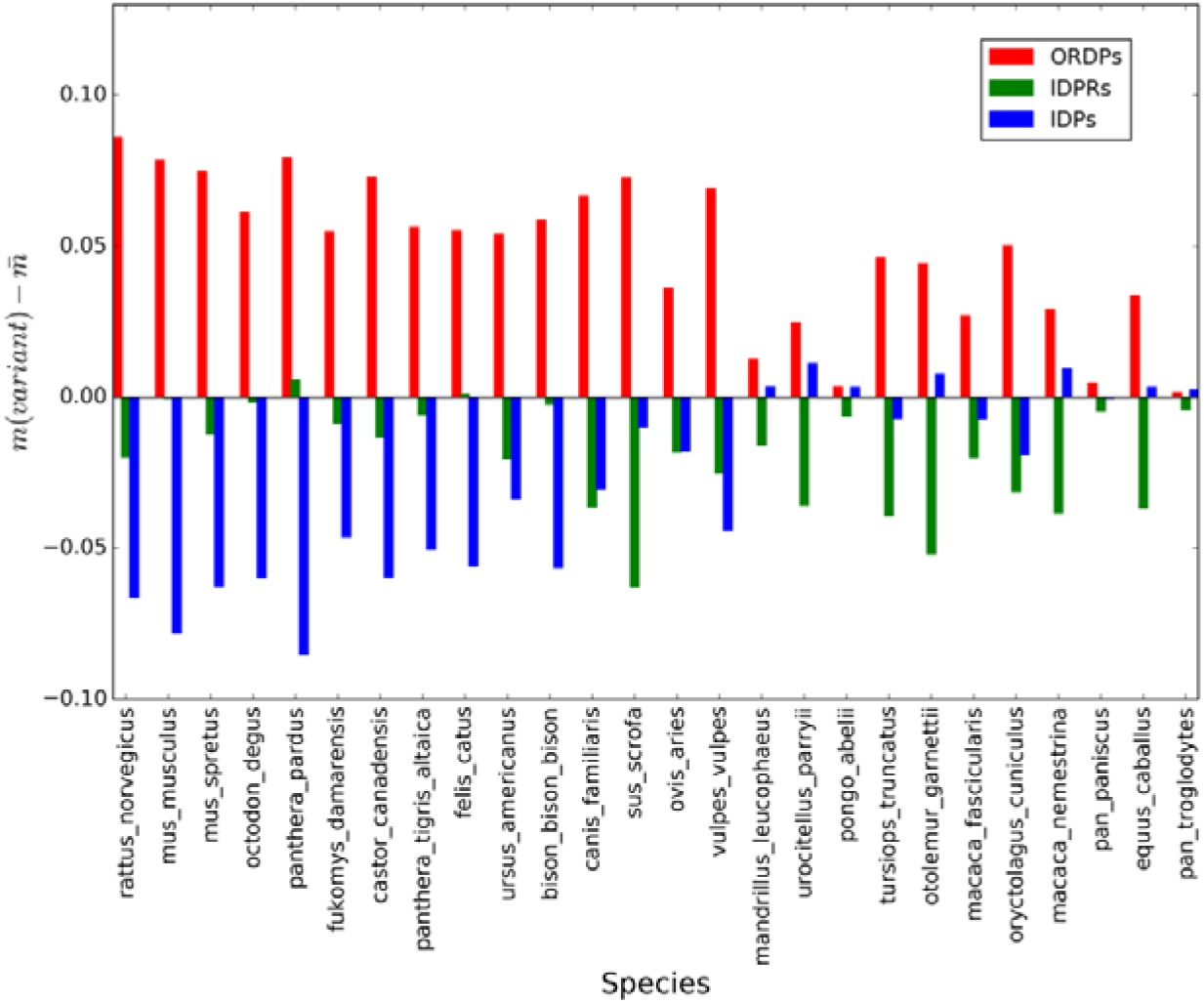
Values of the slopes associated with the variants of disorder, taking Human as the reference.. On the X-axis, we report the 26 eukaryotic species that we considered in the present analysis. Colored bars represent the normalized slopes associated with ORDPs (red), IDPRs (green), and IDPs (blue), with respect to the average values of slopes associated with all variants of disorder. In all comparisons, red bars are the highest, thus suggesting that changes in ORDPs (steeper slopes) play a more powerful role than changes in IDPs and IDPRs (less steep slopes) systematically.

**Figure 3.**
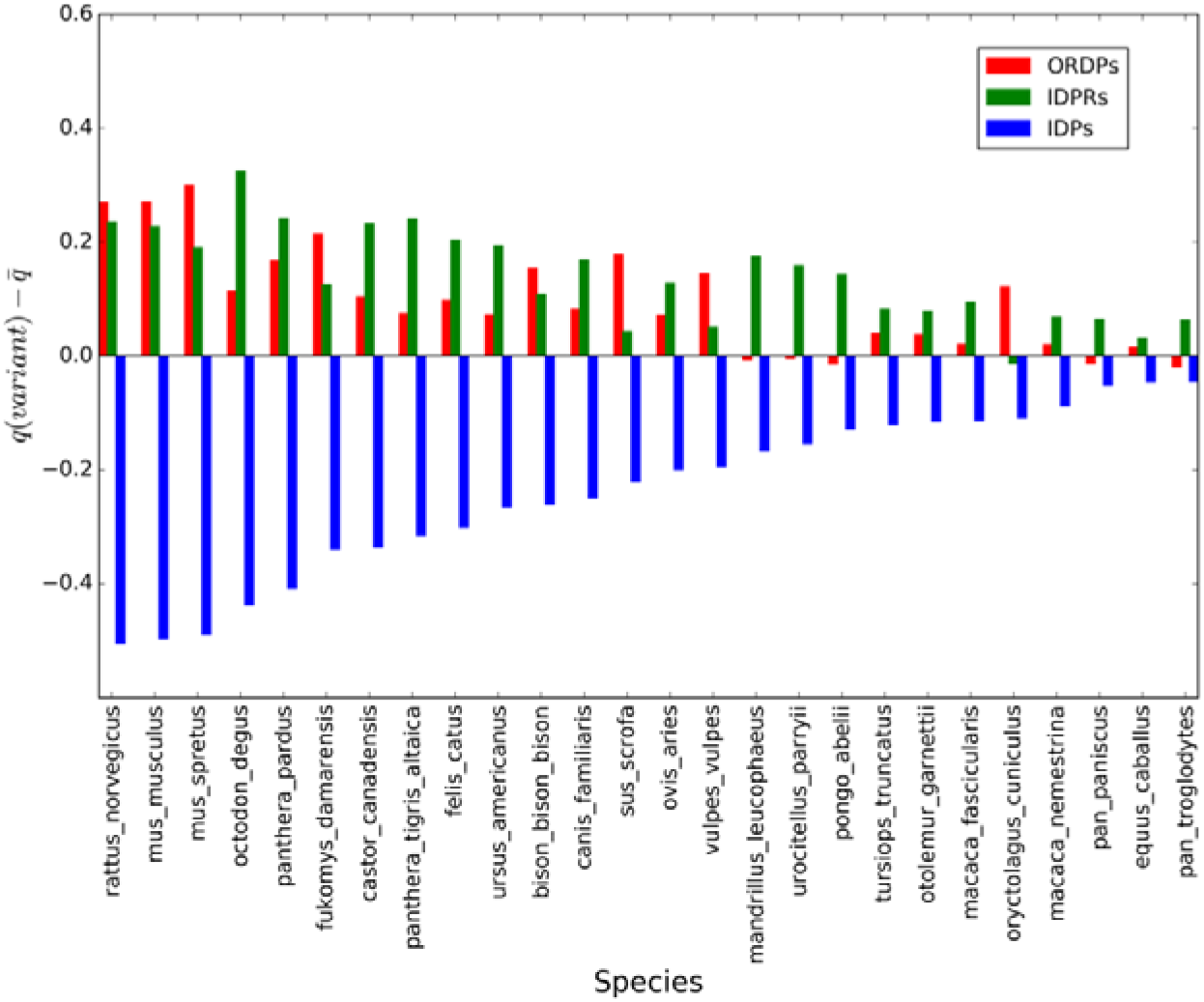
Values of the intercepts associated with the variants of disorder, taking Human as the reference. On the X-axis, we report the 26 eukaryotic species that we considered in the present analysis. Colored bars represent the normalized intercepts associated with ORDPs (red), IDPRs (green), and IDPs (blue), with respect to the average values of intercepts associated with all variants of disorder. In all comparisons, blue bars are the lowest, thus suggesting that IDPs are systematically more reactive at the ‘start’ of the speciation process.

Following the procedure described in Material and Methods, we tested the first null hypothesis that the slopes of the regression lines are all equal. We found that differences between slopes are very significant () in all comparisons, except for *pan troglodytes*, *pongo abelii, equus caballus, urocitellus parryii*, and *ovis aries* (). In these organisms having parallel regression lines, we tested the second null hypothesis that their intercepts are all the same. We found that the intercepts are significantly different () in all comparisons.

As a general trait, IDPs are characterized by the lowest values of the intercepts in all comparisons (blue bars in Fig. 3), thus suggesting that they are systematically more susceptible at the ‘start’ of the speciation process by diverging phenotypically before of ORDPs and IDRPs. On the other hand, ORDPs are characterized by the highest values of the slopes (red bars in Fig. 2) in all comparisons, thus suggesting that they evolve phenotypically faster than IDPRs and IDPs by accumulating nucleic acid changes. It is worth noting that with closely related (allied) species (i.e., *pan troglodytes*, *pongo abelii, mandrillus leucophaeus, macaca nemestrina, macaca fascicularis*, and *pan paniscus)* the fit is good and the points are close to the best-fit line (see Figs S1). In these cases the intercept on the X-axis is near zero (see Fig. 3 and Figs S1). This observation suggests that *genic* differences (i.e., amino acid changes) may have played a significant role in the *pan troglodytes /pongo abelii /pan paniscus – human* divergence (i.e., “genic speciation”), whereas in the other cases *non-genic* differences (both within and external to genes) triggered the separation from the ancestral type into two different species (i.e., “chromosomal speciation”). This is clear by plotting the sum of the square of residuals between the experimental data and the linear fit against the X-intercept values (Fig. 4).

**Figure 4.**
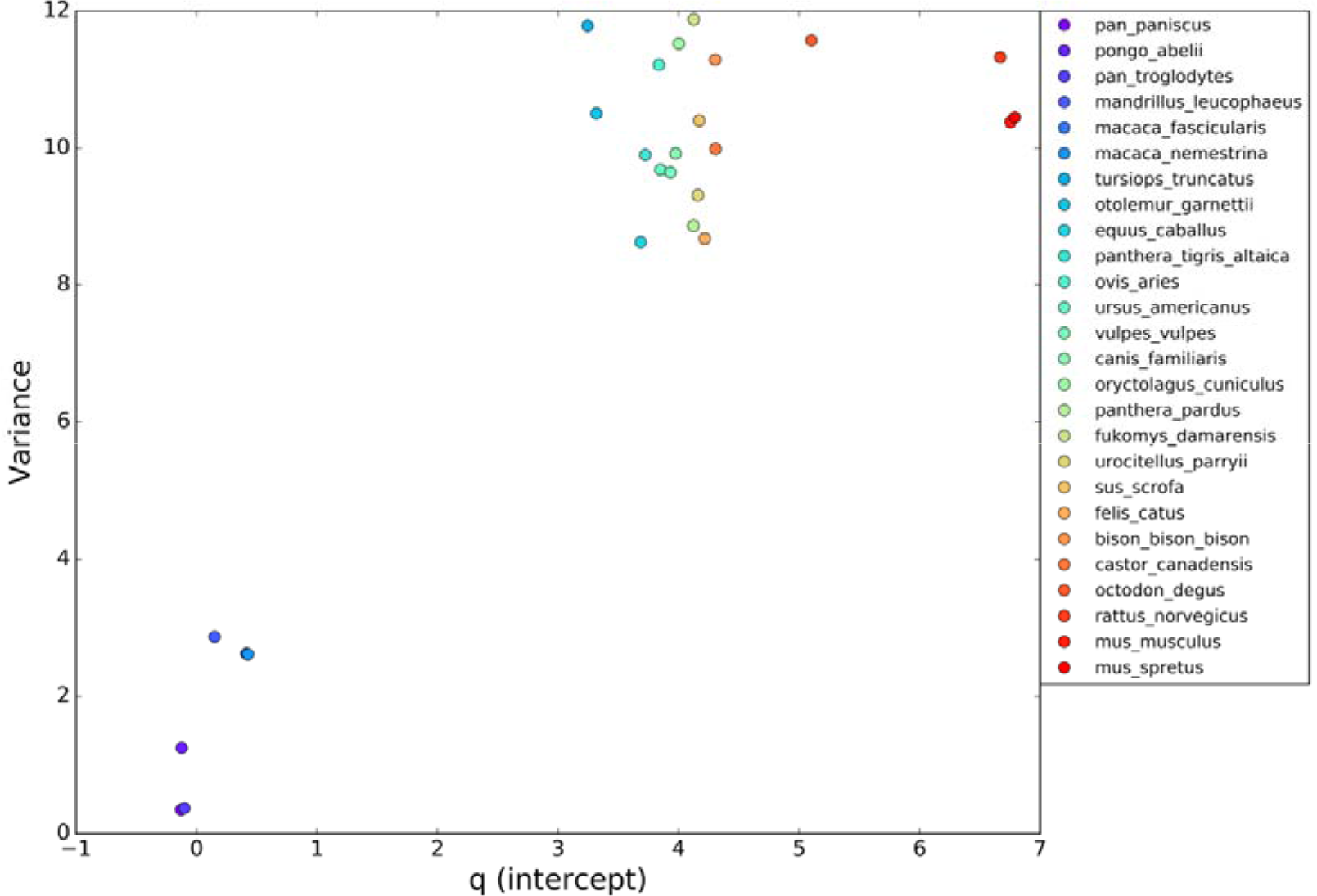
Plot of the sum of the square of residuals between the experimental data and the linear fit against the X-intercept values.

Interestingly, primates are grouped on the bottom-left of the panel, meaning that nucleic acid changes mutations more easily result in amino acid changes mutations, increasing the phenotypic divergence. Conversely, in all the other comparisons on the top-right of Fig. 4, synonymous changes had a chance to accumulate on the coding sequences, leading to a broader variance around the linear-fit.

### 2.3. Similar results are found considering Mus Musculus as the reference

All the results shown above are relative to human. To confirm the general validity of these observations, we repeated the analyses above by considering Mus Musculus as the reference. Thus, we progressively confronted coding sequences in Mus Musculus (separated in ORDPs, IDPRs, and IDPs) with their homologs from 25 eukaryotes. All comparisons produced results similar to Fig. 1 (see Figs S2). Therefore, for each species in contrast with Mus Musculus and each variant of disorder, we retrieve one value of the intercept and one value of the slope. Then, we calculated the normalized values of intercepts and slopes, in the same way as has done in the previous section (Figs. 5 and 6).

**Figure 5.**
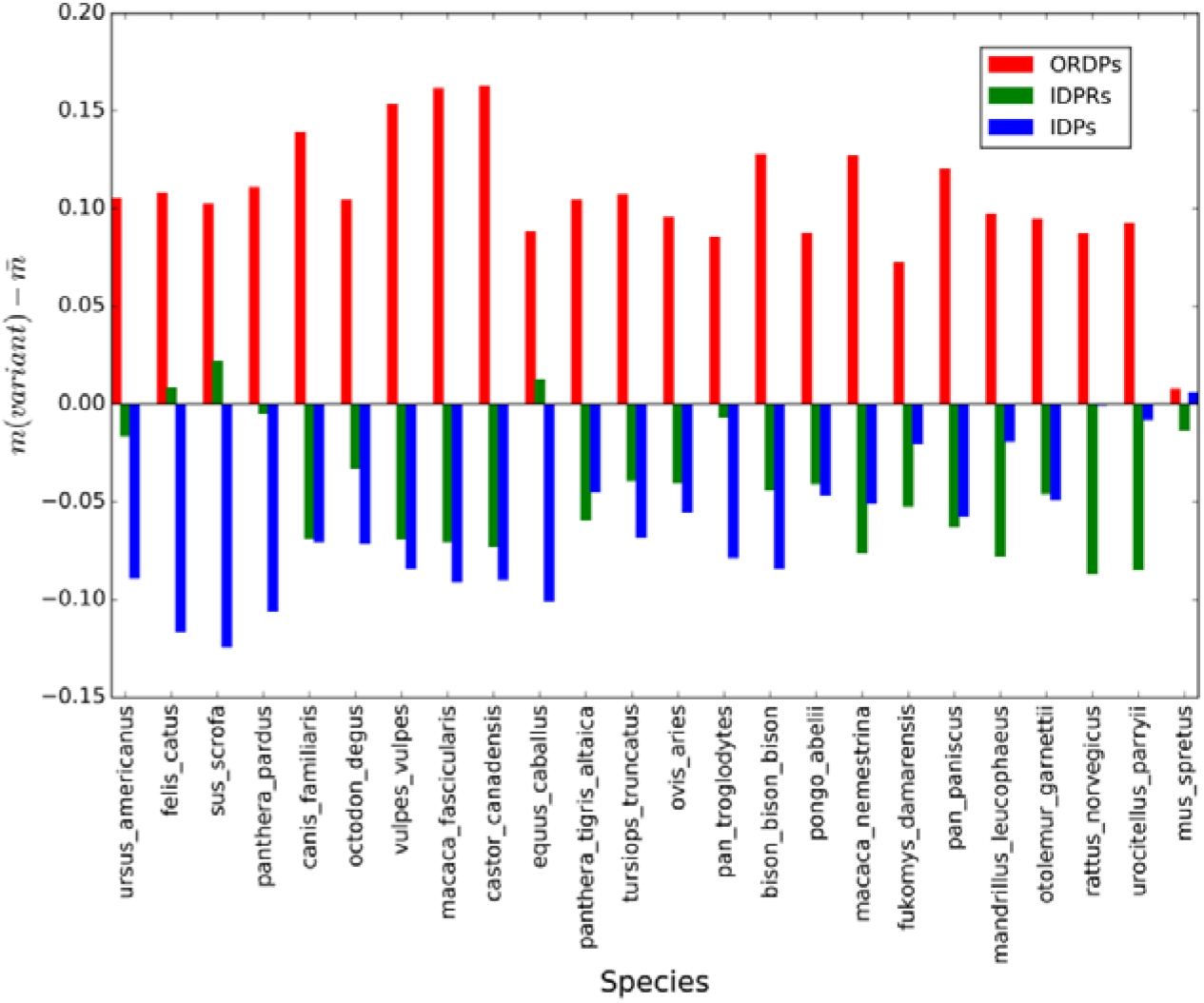
Values of the slopes associated with the variants of disorder, taking Mus Musculus as the reference. On the X-axis, we report the 26 eukaryotic species that we considered in the present analysis. Colored bars represent the normalized slopes associated with ORDPs (red), IDPRs (green), and IDPs (blue), with respect to the average values of slopes associated with all variants of disorder. In all comparisons, red bars are the highest, thus suggesting that changes in ORDPs (steeper slopes) play a more powerful role than changes in IDPs and IDPRs (less steep slopes) systematically.

**Figure 6.**
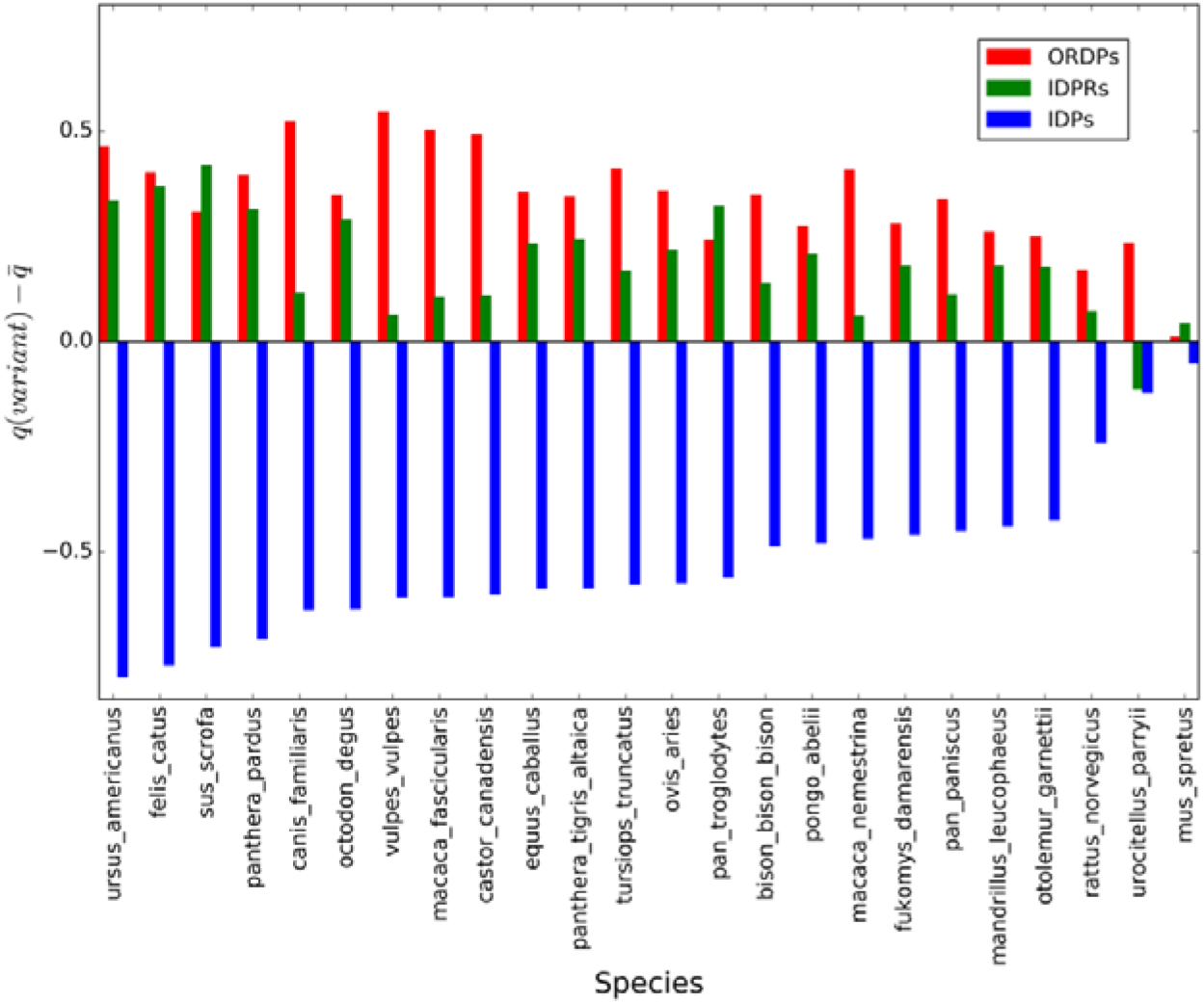
Values of the intercepts associated with the variants of disorder, taking Mus Musculus as the reference. On the X-axis, we report the 26 eukaryotic species that we considered in the present analysis. Colored bars represent the normalized intercepts associated with ORDPs (red), IDPRs (green), and IDPs (blue), with respect to the average values of intercepts associated with all variants of disorder. In all comparisons, blue bars are the lowest, thus suggesting that IDPs are systematically more reactive at the ‘start’ of the speciation process.

Following the procedure described in Material and Methods, we tested the first null hypothesis that the slopes of the regression lines are all equal. We found that differences between slopes are very significant () in all comparisons, except for *mus spretus* (). For this organism, we tested the second null hypothesis that the intercepts of the regression lines are the same. We found that the intercepts are significantly different ().

Similarly to what observed in the previous section, IDPs are characterized by the lowest values of the intercepts (blue bars in Fig. 3), whereas ORDPs are marked by the highest values of the slopes (red bars in Fig. 2). Therefore, we found further evidence that IDPs are more susceptible at the ‘start’ of the speciation process, whereas ORDPs further accelerate the phenotypic divergence once they are recruited.

## 3. Discussion

The main focus of this study is to investigate the role of different variants of disorder in the speciation process. To address this purpose, we separated the human proteome (taken as a reference) in three variants of proteins characterized by different structural and functional properties [12]: i) ordered proteins (ORDPs), ii) mostly ordered proteins with long intrinsically disordered protein regions (IDPRs), and iii) intrinsically disordered proteins (IDPs).

Motivated by Wolfe and Sharp [14], we compared the DNA divergence with the corresponding protein divergence, by confronting human coding sequences (separated in ORDPs, IDPRs, and IDPs) and their homologues sequences from 26 eukaryotes.

The intercepts on the X-axis are evidence that a broad nucleic acid level process, rather than a specific protein level process, initiates speciation in most cases [15]. Beyond that, our results provide evidence that particular variants of disorder will contribute before others. In particular, IDP-encoding genes appear to be less able to accommodate nucleic acid changes at the ‘start’ of the speciation process. Thus, IDPs seem to diverge phenotypically before of ORDPs and IDPRs, potentially triggering the speciation process. Then, once ORDPs start to differ in amino acids, they become phenotypically more reactive to nucleic acid changes than IDPRs and IDPs. Furthermore, our findings show that, once the phenotypic divergence has started, IDP-encoding genes become freer to accept nucleic acid changes and especially synonymous mutations, which are generally assumed to be neutral. This observation confirms the central role of IDPs for protein evolvability because neutral mutations usually leave the primary function of the molecule unchanged while paving the way for new features to emerge [6].

Noteworthy, similar results are found by considering Mus Musculus as the reference, thus confirming the general validity of our considerations. Here too, IDPs appear to be involved in the early stages of the speciation process, whereas ORDPs appear to accelerate the further phenotypic divergence.

Should IDPs themselves be involved in the early stages of the speciation process, this could be connected to their functional spectra, mainly related to nucleic acid binding and transcription factors (Deiana et al. 2019). Primary changes at the nucleic acid level (GC-pressure mutating both coding and non-coding DNAs at different rates) creates two potentially divergent lines [14]. Rates of mutations in functional regions are expected to be less than the rates of mutations of third codon positions, introns, and intergenic DNA. However, exceptions may be genes encoding nucleic acid-binding proteins, which may need to bind optimally their substrate that progressively differs in GC content [18]. Thus, the mutation rate in nucleic acid binding proteins, which are enriched in IDPs [12], could be higher than in most other proteins. In turn, the higher rate of mutations in some of these genes (e.g., those involved in DNA repair) might lead to directional changes in genomic GC content, resulting in differentiation of species [18]. Similarly, several evolutionary and molecular features have been revealed that make transcription factors, especially the family of KRAB-ZNF transcription factors, strong candidates to play an important role in speciation [10].

In sum, this study provides compelling evidence about the differential roles of different structural variants of proteins in the speciation process, identifying IDPs as a trigger and ORDPs as a further prompt of the phenotypic divergence.

## 4. Methods

### 4.1. Data sources

The proteomes of Homo Sapiens and Mus Musculus were downloaded from the UniProtKB/SwissProt database (manually annotated and reviewed section – https://www.uniprot.org/uniprot/?query=reviewed:yes) [19]. Coding DNA Sequences (CDSs) of Human and Mus Musculus were retrieved by Ensembl Genome Browser 94 (https://www.ensembl.org/index.html) [20]. Only genes with UniProtKB/SwissProt ID have been included to make sure we only consider coding sequences for proteins. We consider only CDSs that start with the start codon (AUG), end with a stop codon (UAG, UAA, or UGA), and have a multiple length of three. Each CDS was translated in the corresponding amino acid sequence and then we filter all sequences that do not have a complete correspondence with a protein sequence in UniProtKB/SwissProt. Incomplete and duplicated gene, sequences with internal gaps, unidentified nucleotides were removed from the analysis. Two lists of 18150 and 14757 CDSs were generated for Human and Mus Musculus, respectively.

### 4.2. Disorder prediction

We identified disordered residues in the protein sequences using MobiDB3.0 (http://mobidb.bio.unipd.it) [21], a consensus database that combines experimental data (especially from X-ray crystallography, NMR, and cryo-EM), manually curated data, and disorder predictions based on various methods.

### 4.3. Classification of the human proteome

The proteomes of Homo Sapiens and Mus Musculus were partitioned into variants of disorder following the operational classification by Deiana et al. [12]. In line with that, we distinguish three variants of human proteins having different structural, functional, and evolutionary properties: *i) ordered proteins* (ORDPs): they have less than 30% of disordered residues, no C- or N-terminal segments longer than 30 consecutive disordered residues as well as no segments longer than 40 consecutive disordered residues in positions distinct from the N- and C-terminus; *ii) proteins with intrinsically disordered regions* (IDPRs): they have less than 30% of disordered residues in the polypeptide chain and at least either one C- or N-terminal segment longer than 30 consecutive disordered residues or one segment longer than 40 consecutive disordered residues in positions distinct from the N- and C-terminus; *iii) intrinsically disordered proteins* (IDPs): they have more than 30% of disordered residues in the polypeptide chain.

### 4.4. Analyses of DNA and protein divergence among homologous sequences

The human proteome (separated in ORDPs, IDPRs, and IDPs) was taken as a reference. For each human gene, we used the Ensembl REST API (option ‘homology/id/:id’) to retrieve its alignment with a homologous sequence in another eukaryotic organism. Then, we compared base substitutions in the coding sequences with amino acid substitutions in the corresponding protein sequences. Thus, for each pair of sequences in comparison, we calculate the percentage of DNA sequence divergence (*DNA div* (%)) as follows:

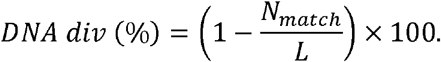

where *N_match_* is the number of matches and *L* is the length of the DNA alignment. The protein sequences were then aligned using the DNA alignments as templates. Then, we calculate the percentage of protein sequence divergence (*Protein div* (%)) as follows:

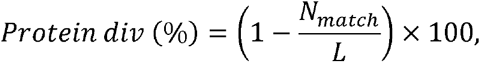

where *N_match_* is the number of matches and *L* is the length of the protein alignment.

### 4.5. Statistical test for comparing multiple linear regressions

For each species in comparison with Homo Sapiens or Mus Musculus, the first step is to compute the regression line between Protein vs. DNA sequence divergence for each variant of disorder. In this way, we can study how nucleic acid changes in each variant are reflected in protein divergence by estimating the slope and the intercept of the regression line.

Next, we test whether the linear regressions associated with the variants of disorder are significantly different by using the protocol defined by Armitage et al. to compare more than two linear regressions [22].

The first null hypothesis is that the slopes of the regression lines are all equal. If the `p – value` is lower than 0.05 then the null hypothesis that the regression lines are parallel is discarded. If the slopes are different, the regression lines cross each other somewhere, and one of the variants has higher *protein* divergence in one part of the plot and lower *protein* divergence in another part of the plot. In this case, there is no point in comparing the intercepts [23]. Indeed, to test the difference between intercepts of two linear regressions with the x-axis, it is sufficient to calculate their interception point and show that its ordinate is different from 0.

Alternatively, if we accept the null hypothesis that the regression lines are parallel, we test the second null hypothesis that their intercepts are all the same. If the `p – value` is lower than 0.05, we conclude that the lines are distinct but parallel. Otherwise, if `p – value > 0.05` then there is no evidence that the regression lines are different.

## Acknowledgements

The authors warmly thank prof. D. R. Forsdyke for the critical reading of the manuscript and for his generous share of ideas, constructive comments and suggestions. We would also like to thank Dr. Marco Punta of ICR London, for his valuable discussions.

## Author contributions

Forcelloni conceived the study. Forcelloni conducted the analyses. Forcelloni and Giansanti wrote the manuscript. Both authors read and approved the final manuscript.

