## Supplementary material for "Mutations in disordered proteins as early indicators of nucleic acid changes triggering speciation": Figs S1

**Supplementary information**

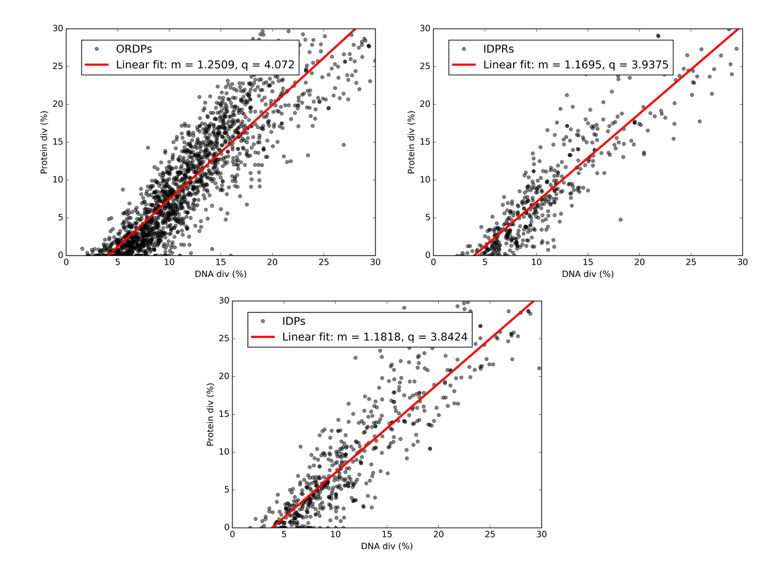

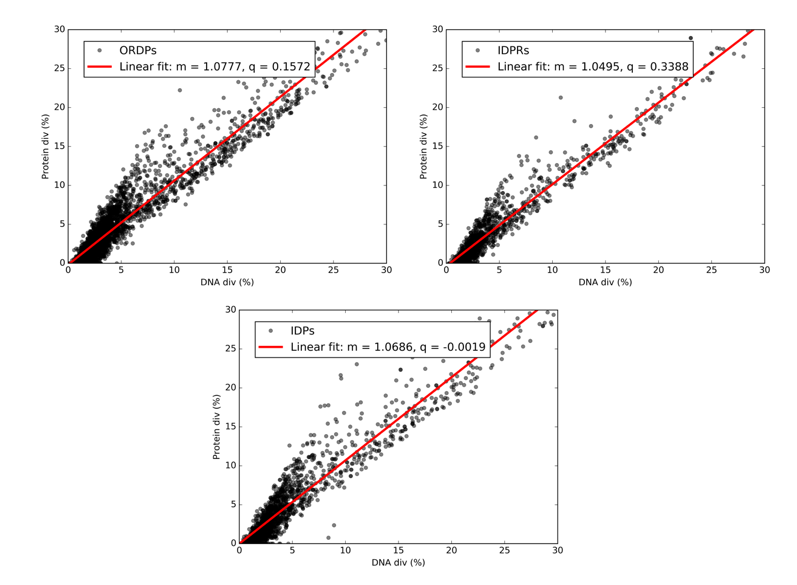

Oryctolagus cuniculus	Mandrillus leucophaeus

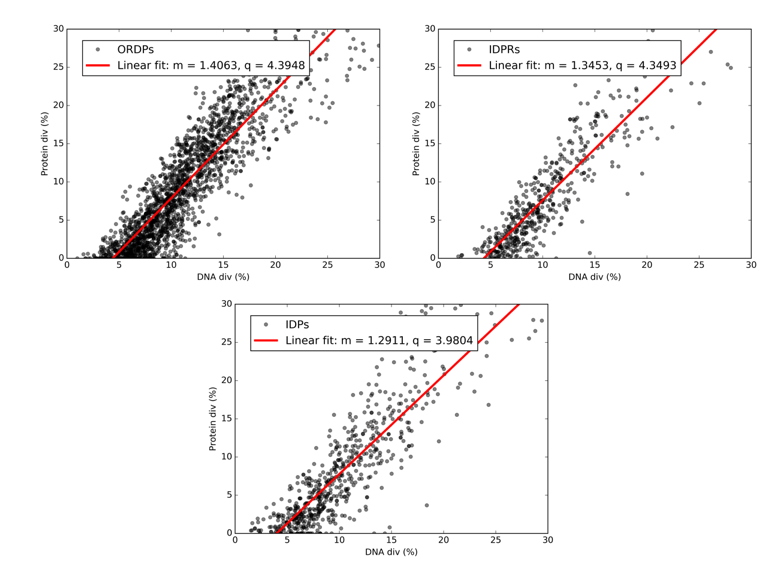

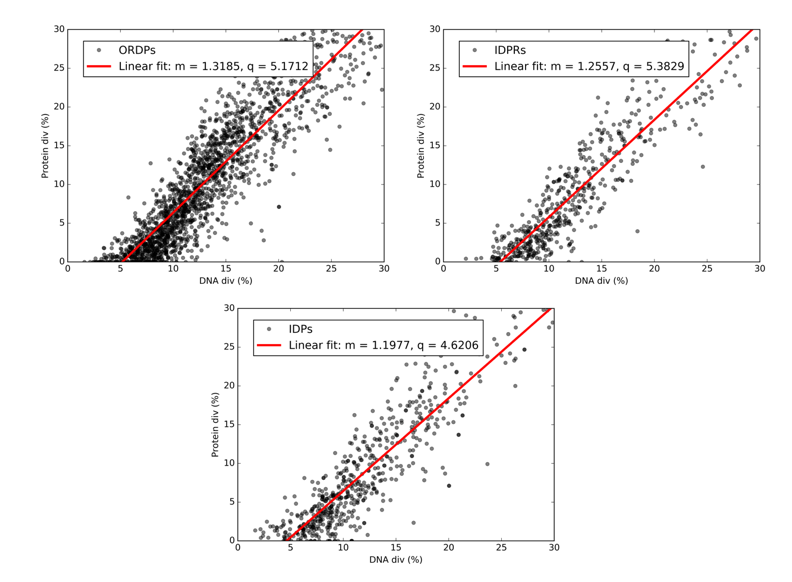
Bison bison bison	Octodon degus

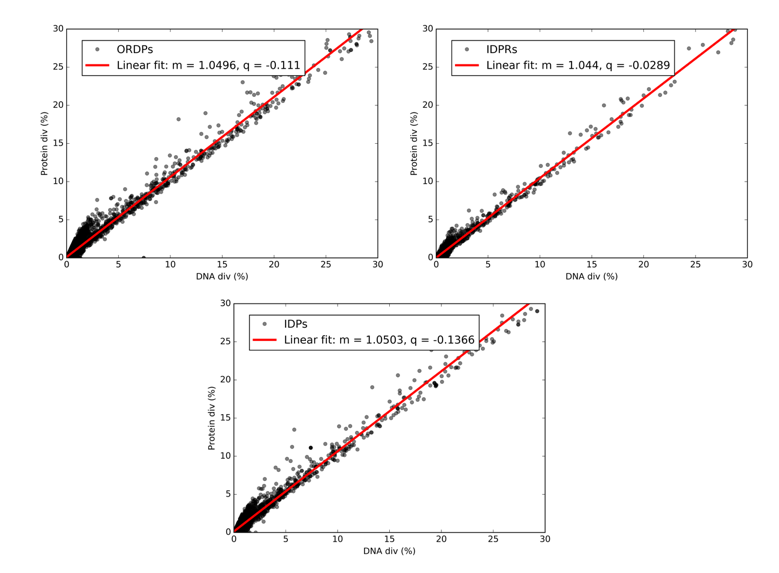

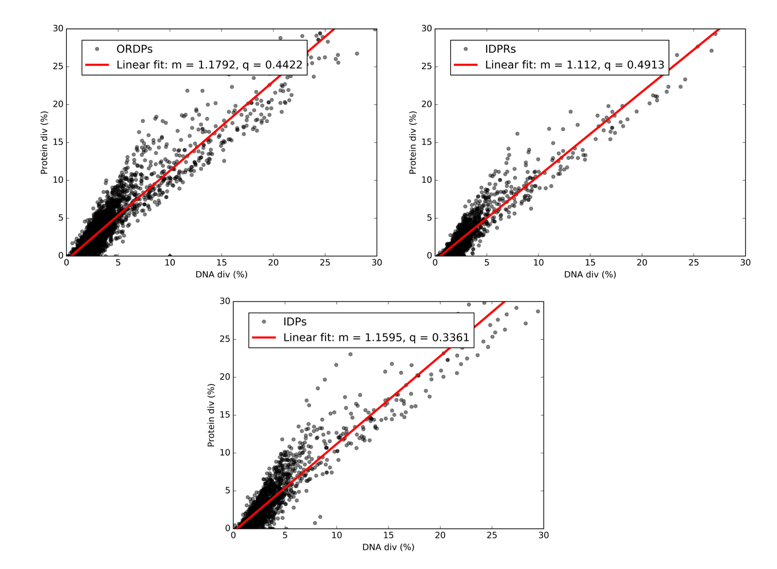

Pan troglodytes	Macaca nemestrina

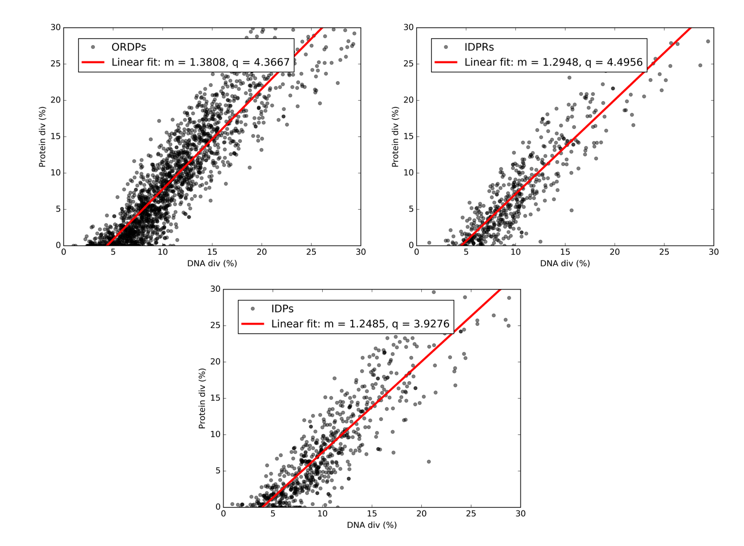

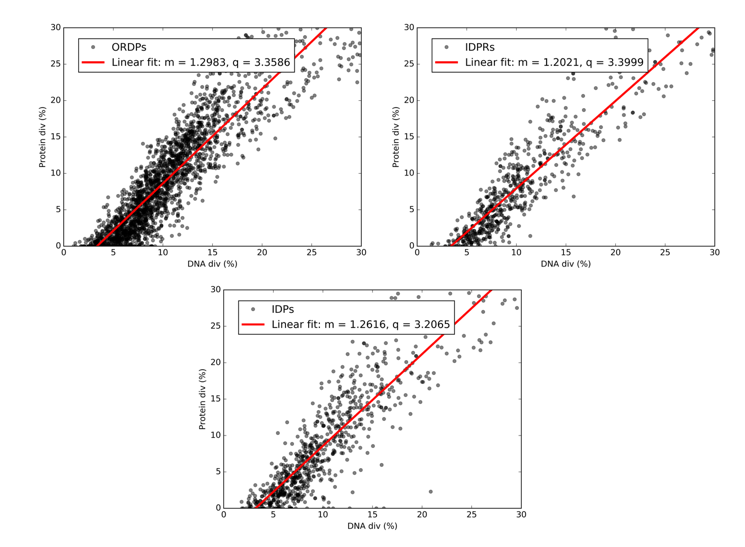

Castor canadensis	Otolemur garnettii

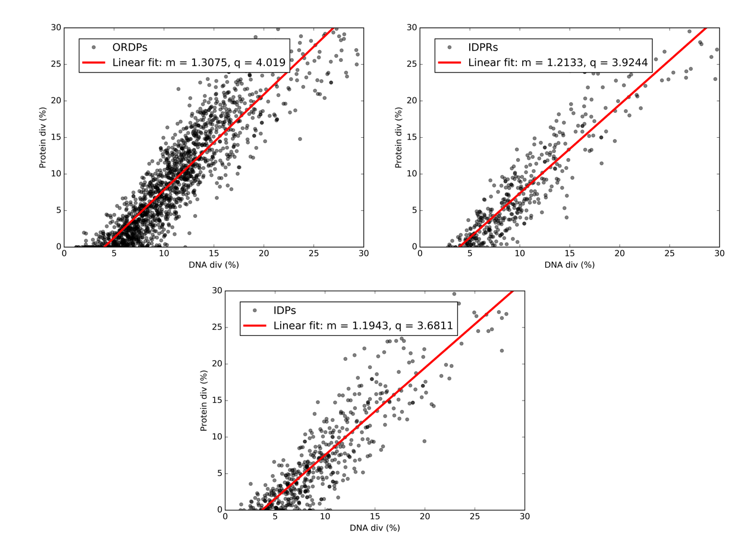

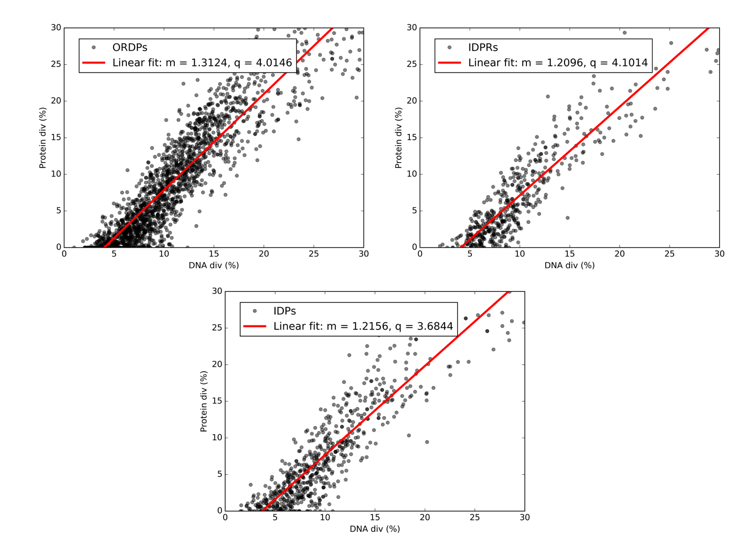

Vulpes vulpes	Canis familiaris

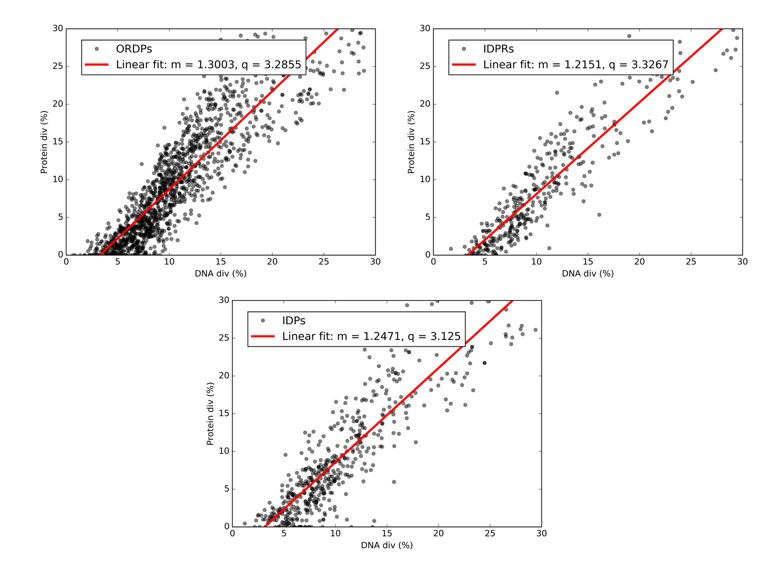

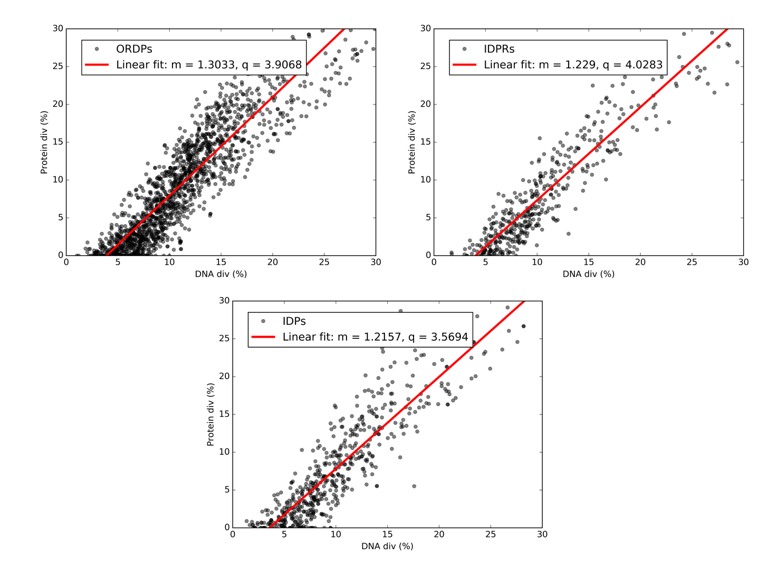

Tursiops truncates	Ursus americanus

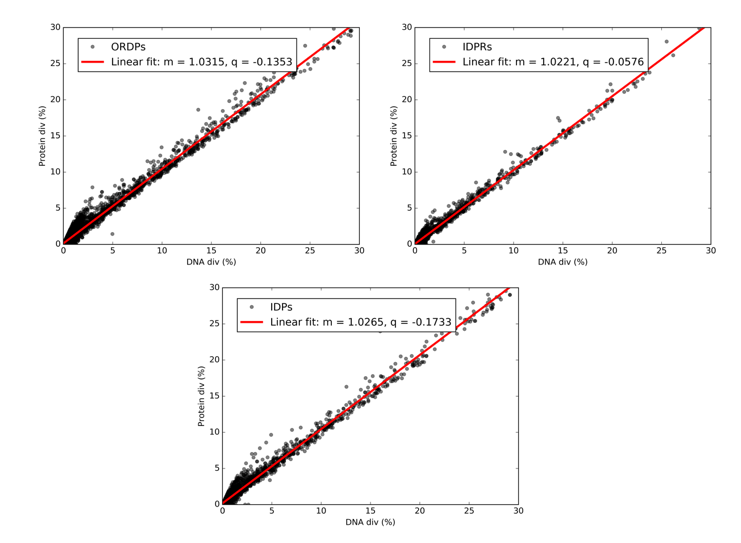

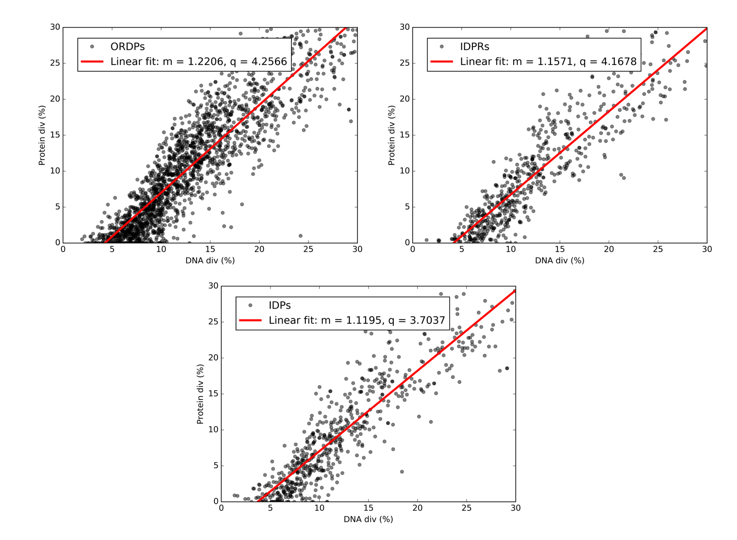

Pan paniscus	Fukomys damarensis

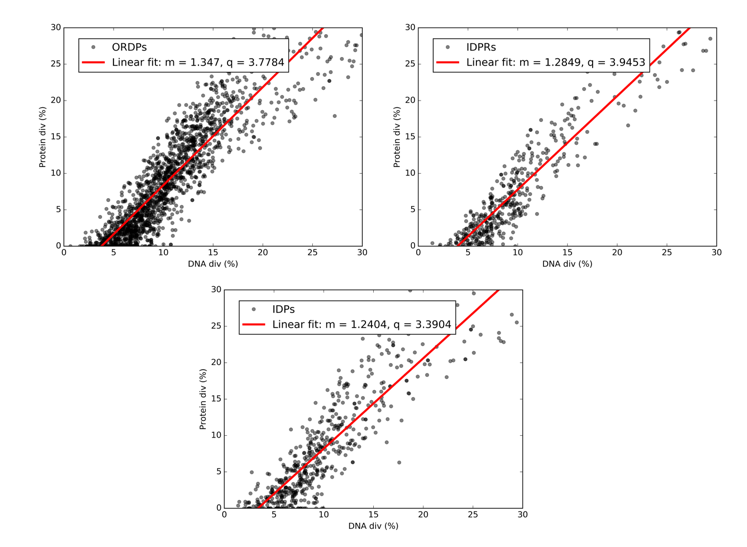

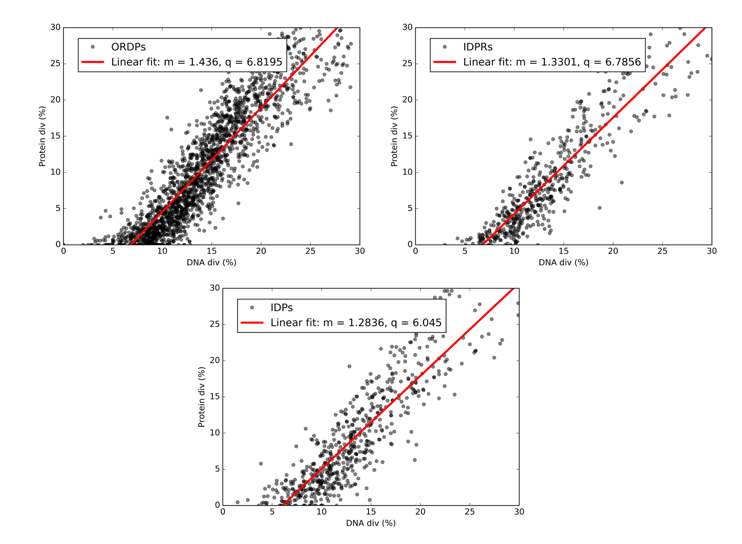

Panthera tigris altaica	Rattus norvegicus

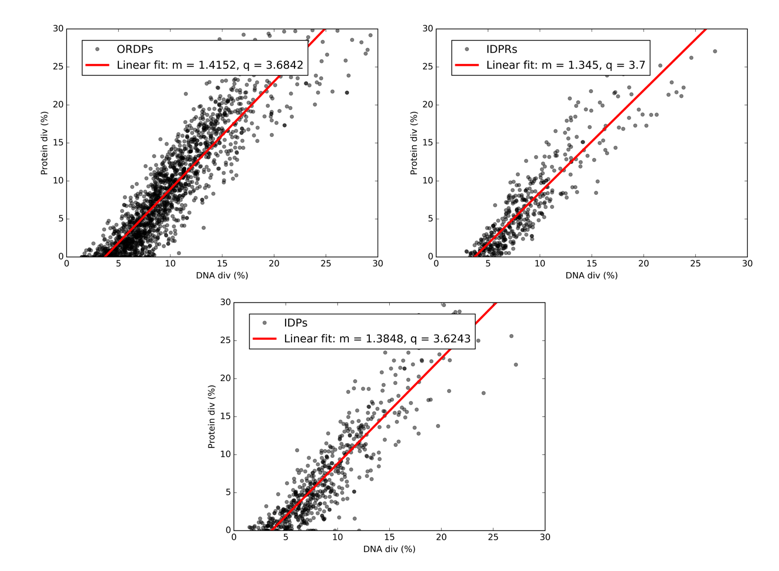

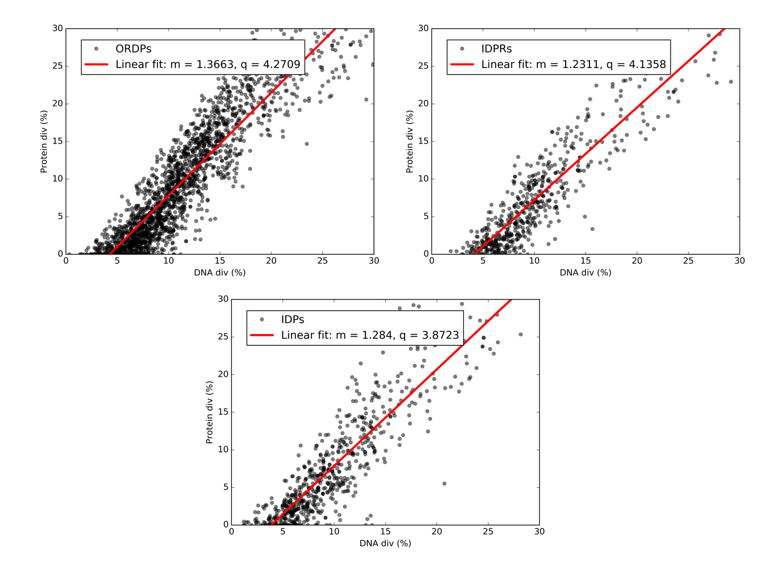

Equus caballus	Sus scrofa

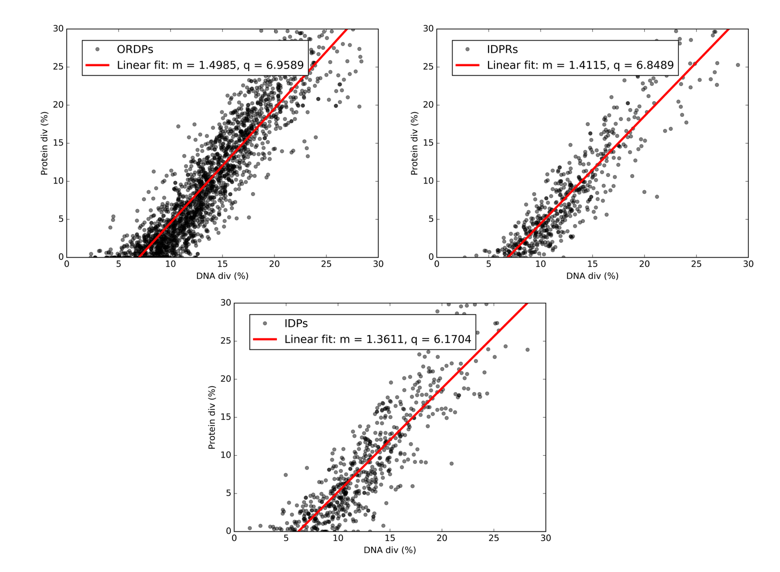

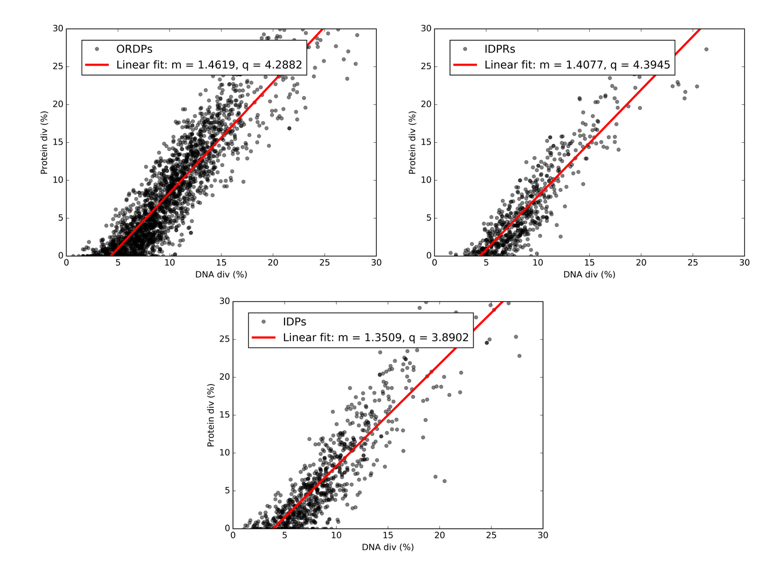

Mus spretus	Felis catus

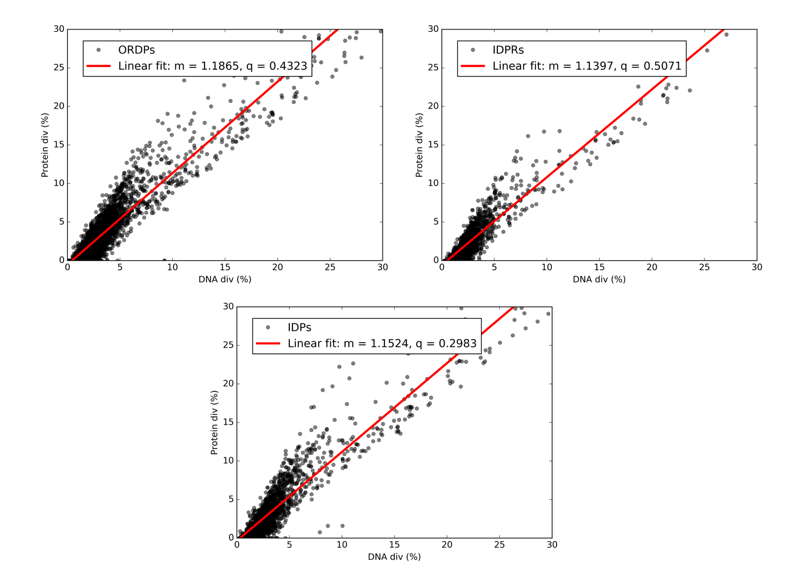

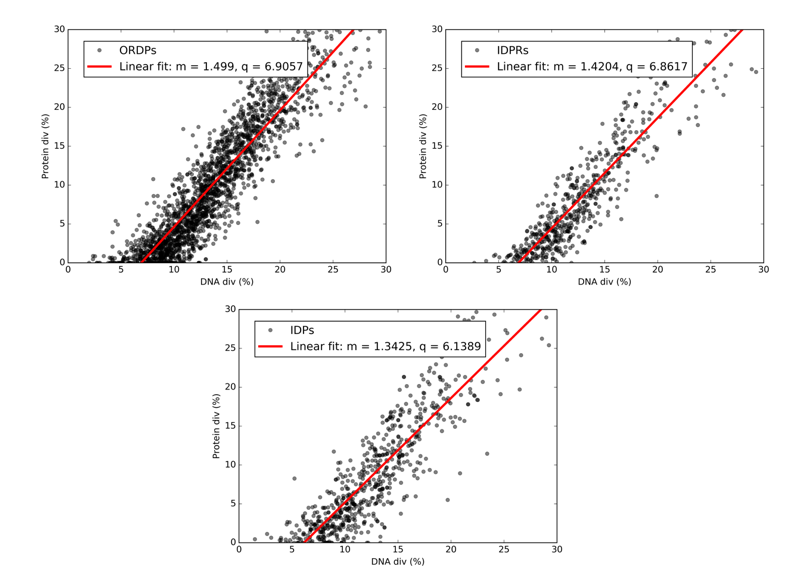

Macaca fascicularis	Mus musculus

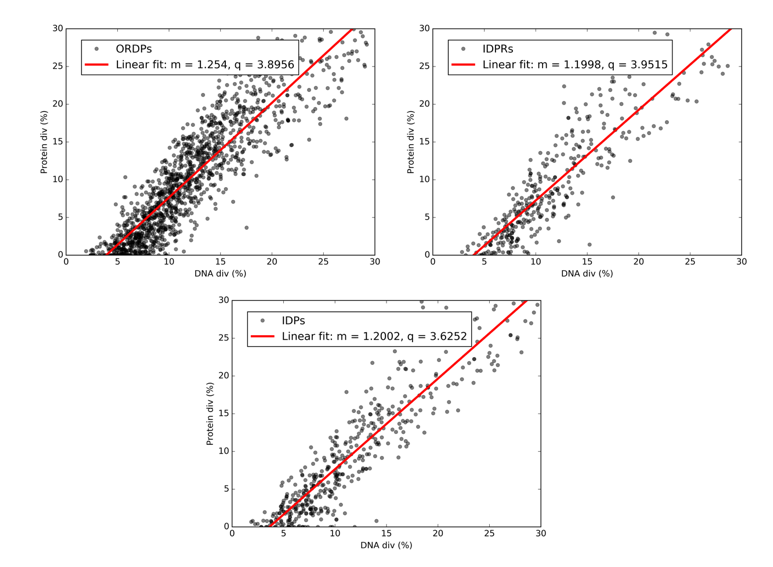

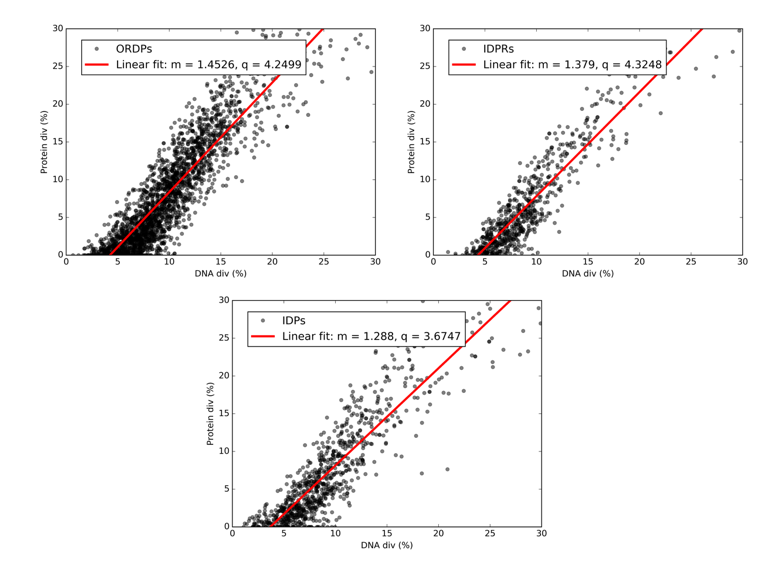

Ovis aries	Panthera pardus

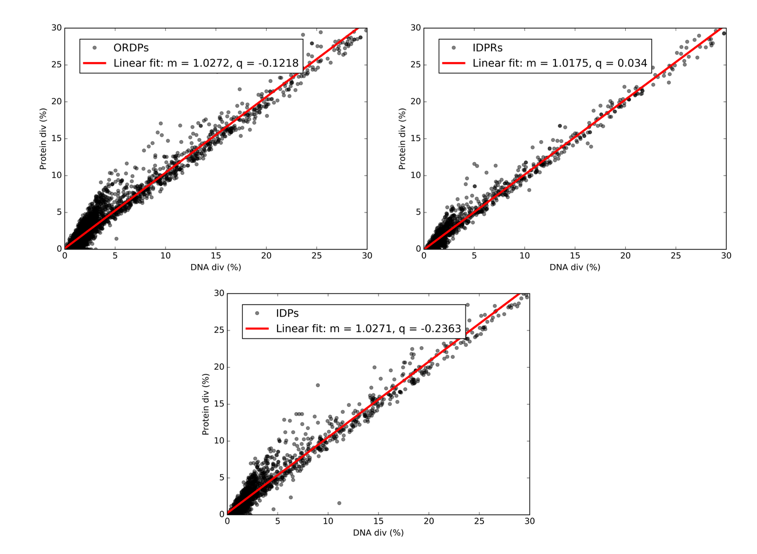

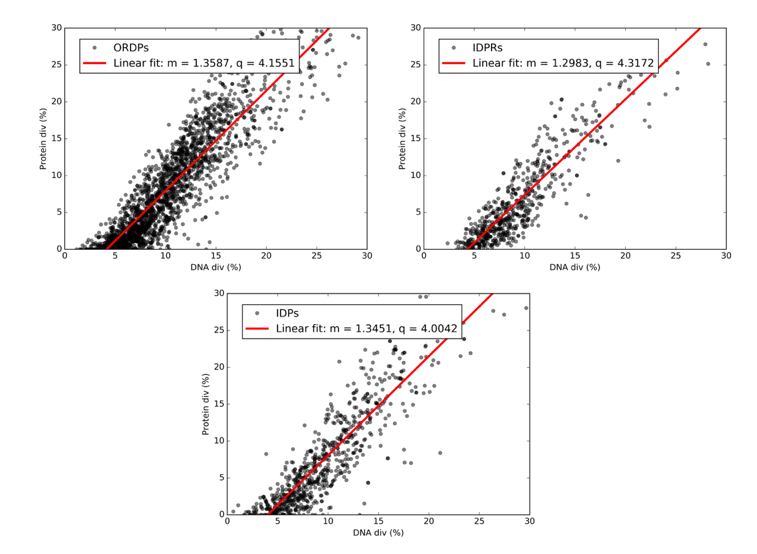

Pongo abelii	Urocitellus parryii

**Figure S1: DNA divergence Vs. Protein divergence plots.** Relationship between nucleotide (DNA div) and amino acid (Protein div) sequence divergence obtained by confronting progressively human coding sequences (separated in ORDPs, IDPRs, and IDPs) with their homologs from 26 eukaryotes. Each point corresponds to an individual gene. In each panel, we report the best-fit line, together with the associated values of the slope (m) and the intercept (q) in the legend.

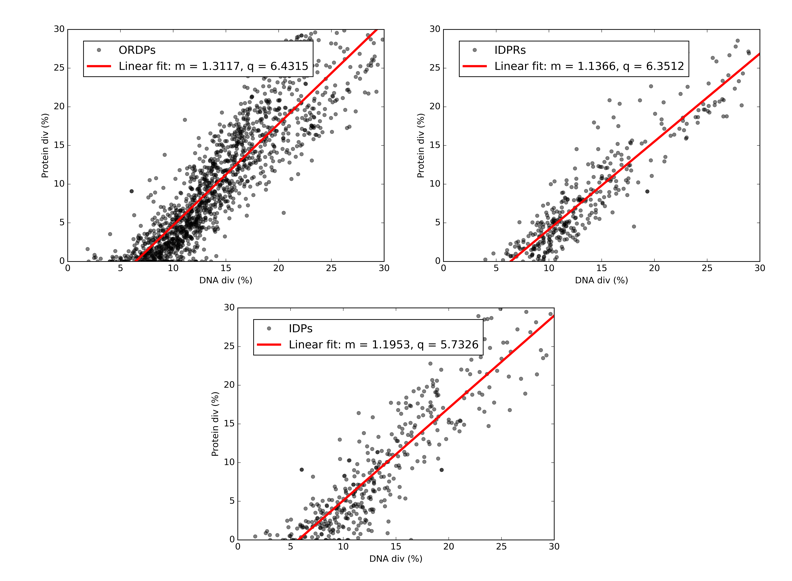

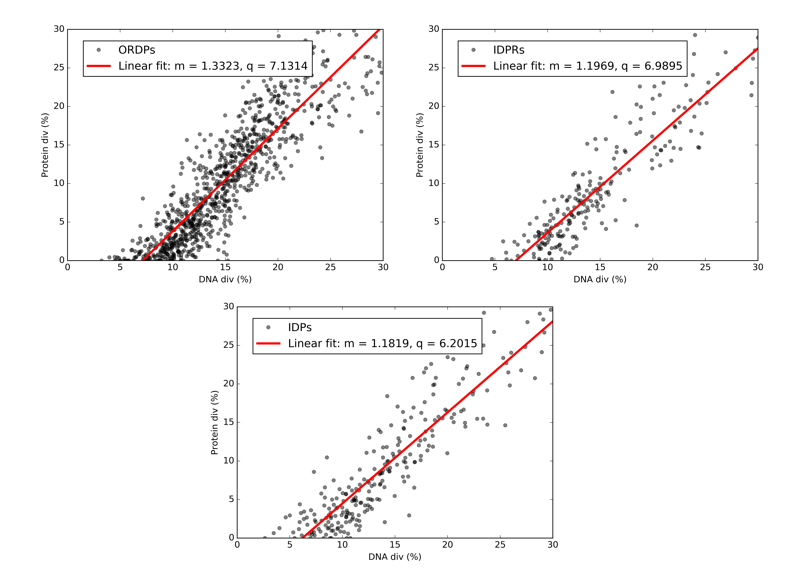

Mandrillus leucophaeus	Ovis aries

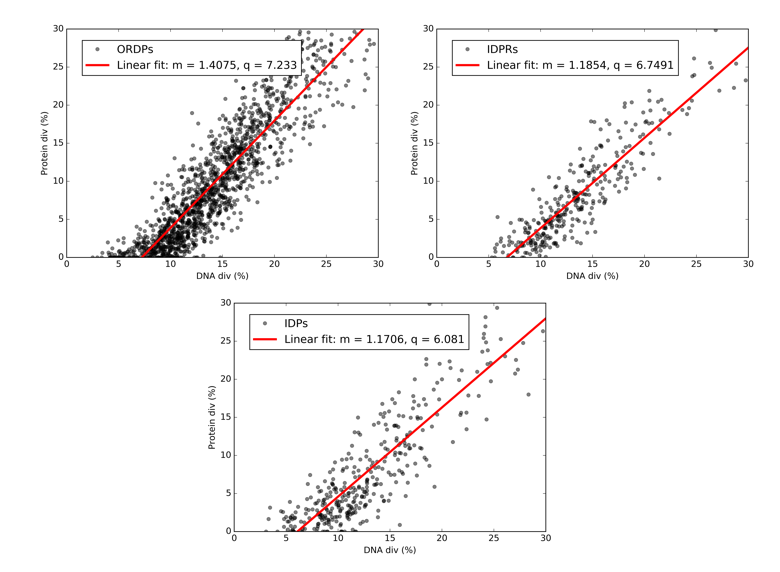

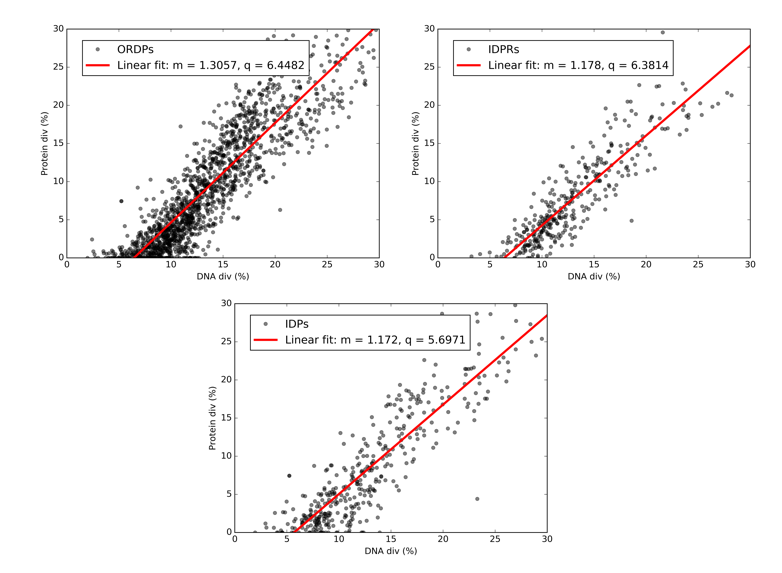

Vulpes Vulpes	Pongo abelii

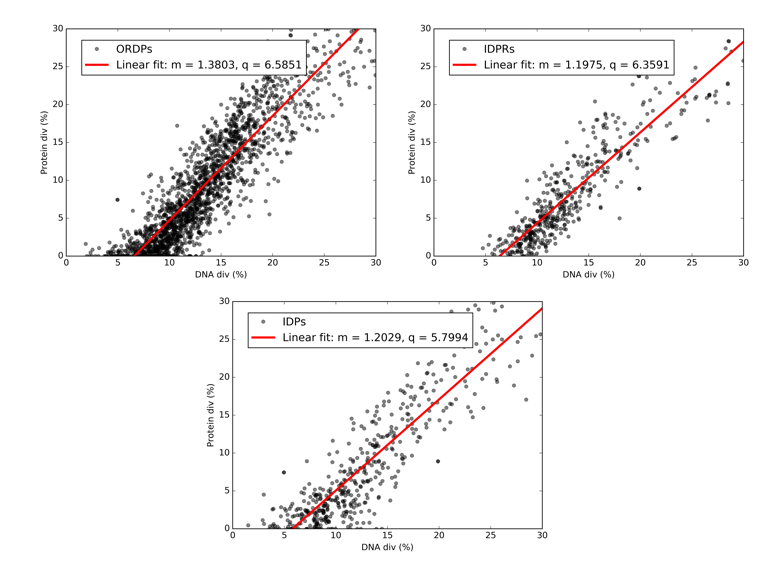

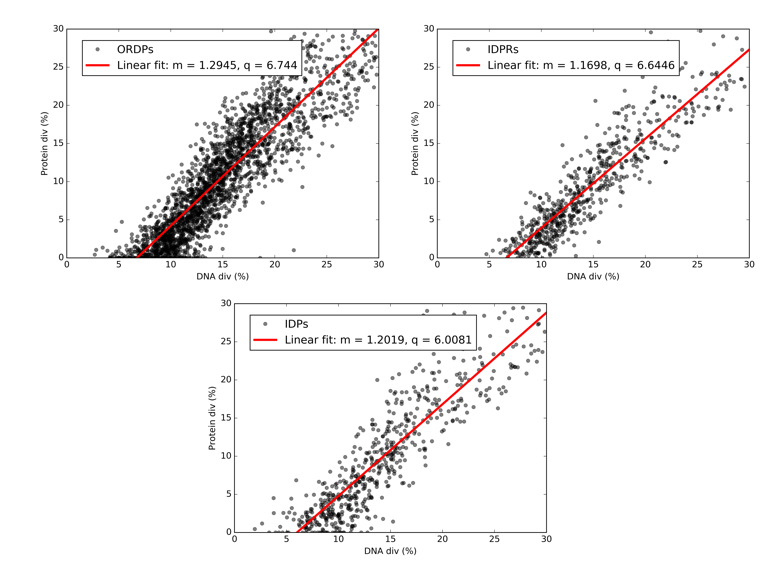

Pan paniscus	Fukomys damarensis

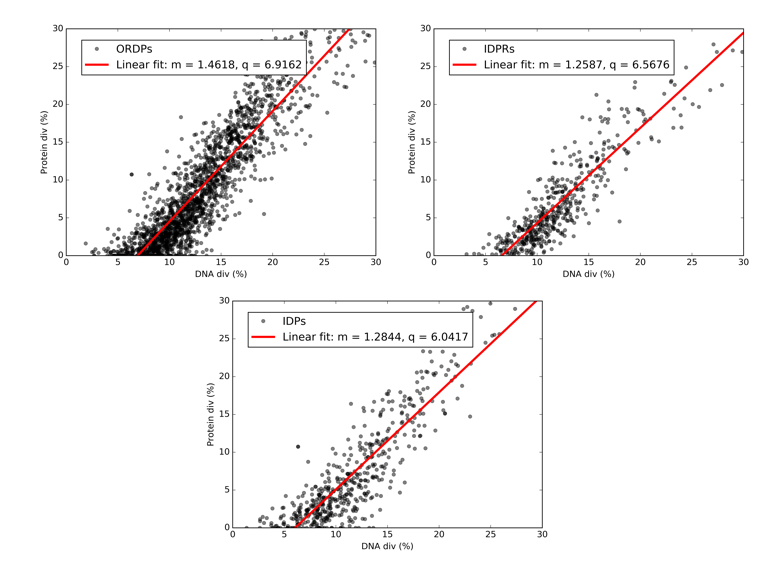

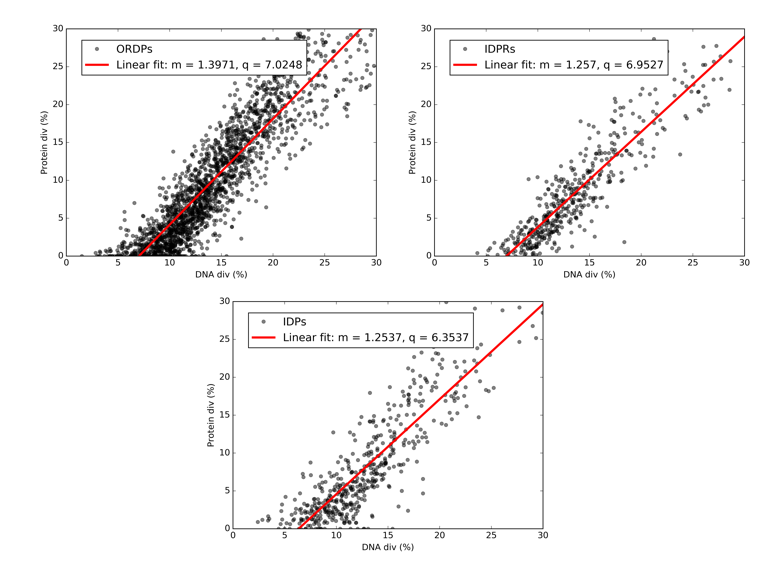

Macaca nemestrina	Otolemur garnettii

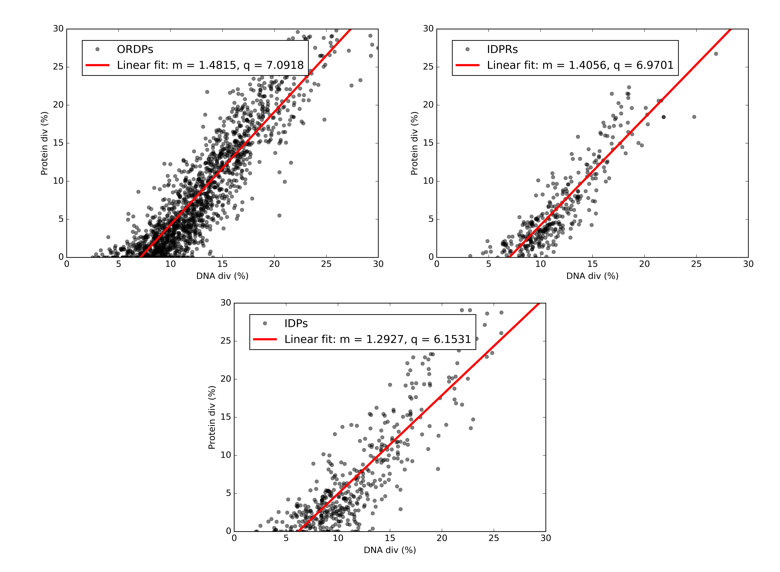

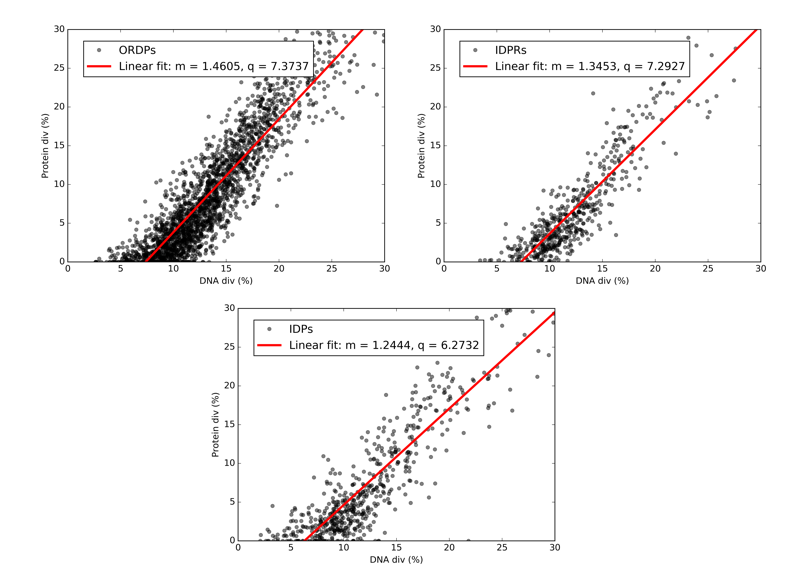

Equus caballus	Panthera pardus

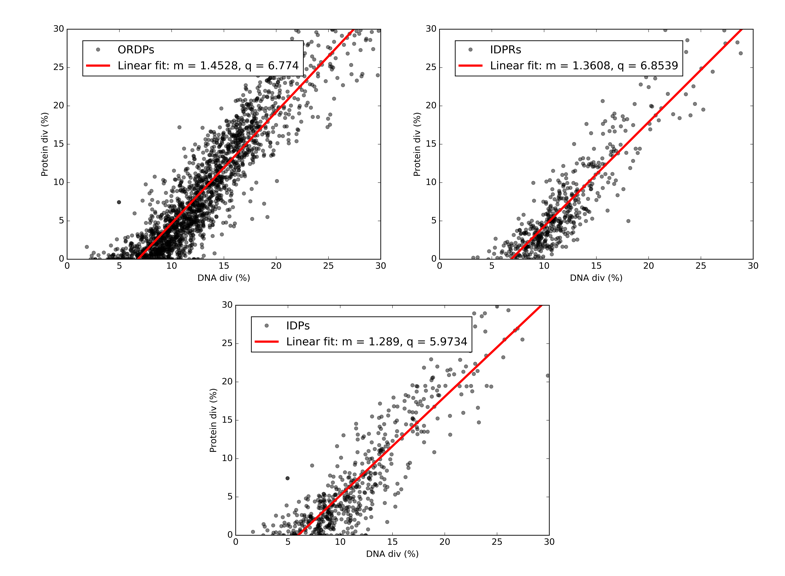

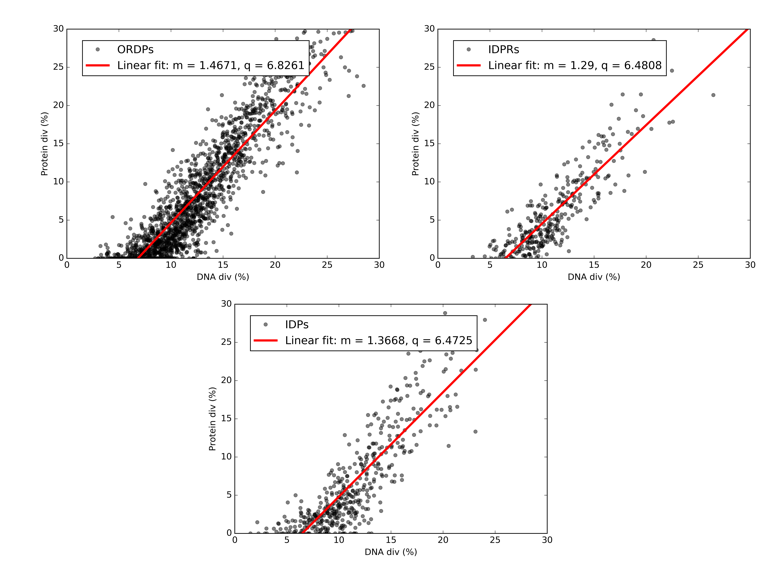

Pan troglodytes	Urocitellus parryii

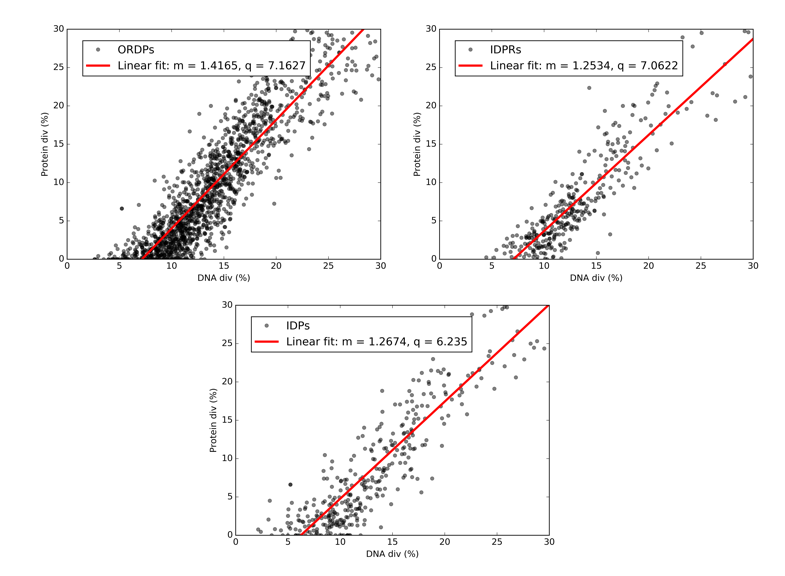

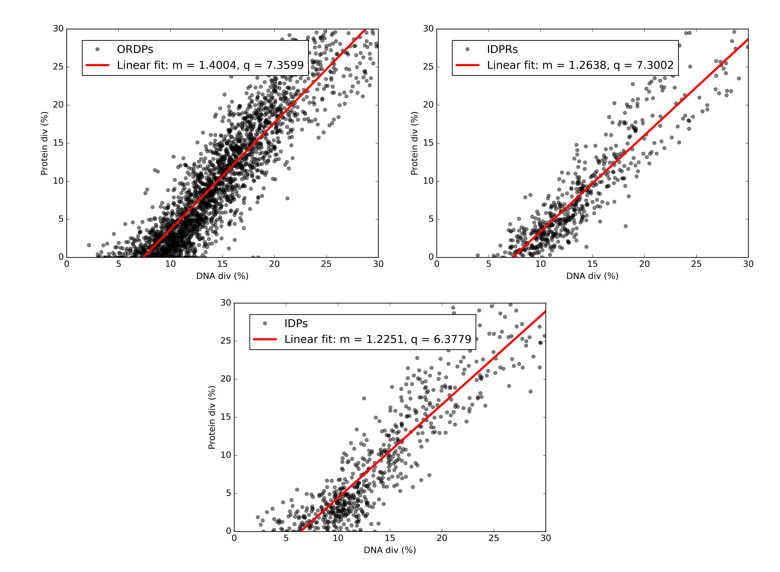

Panthera tigris altaica	Octodon degus

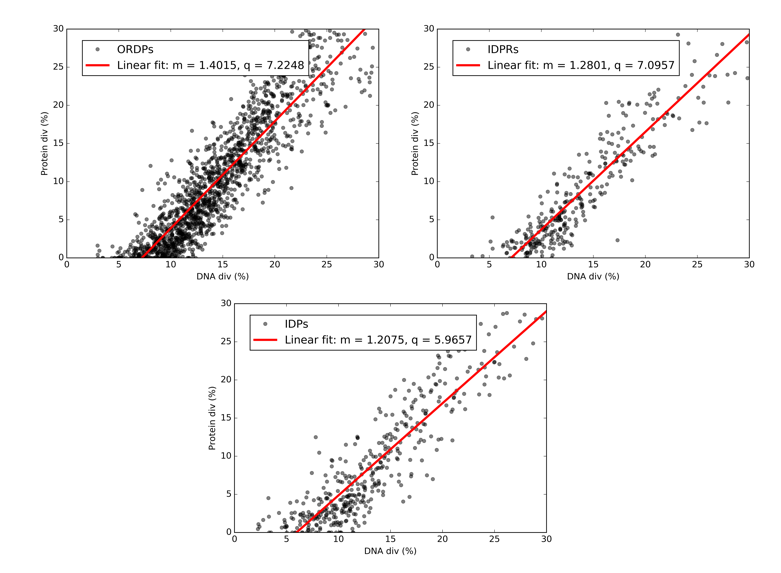

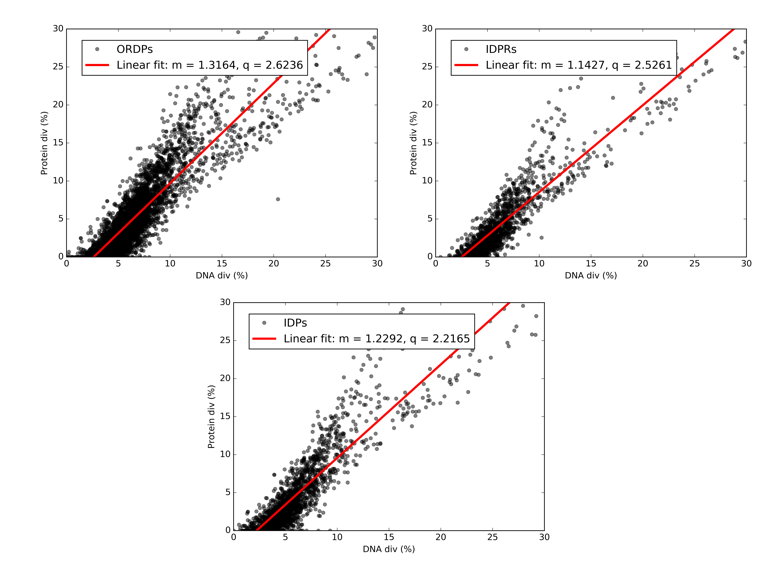

Ursus americanus	Rattus norvegicus

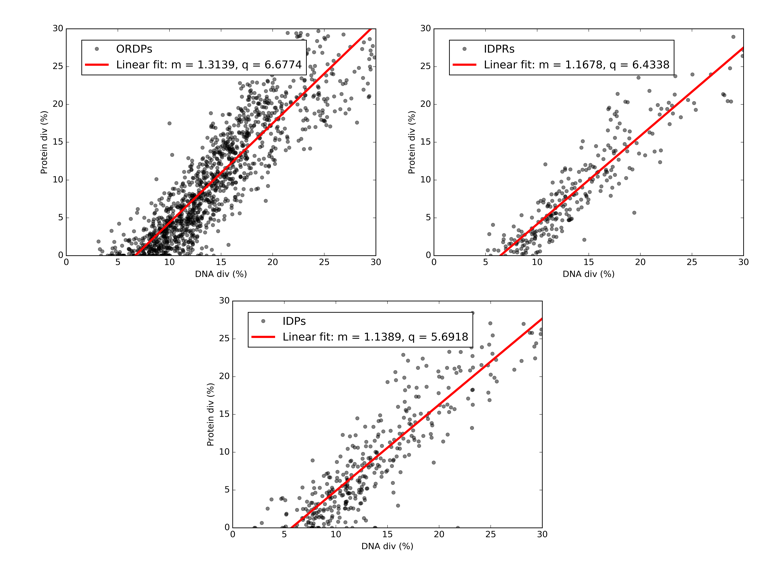

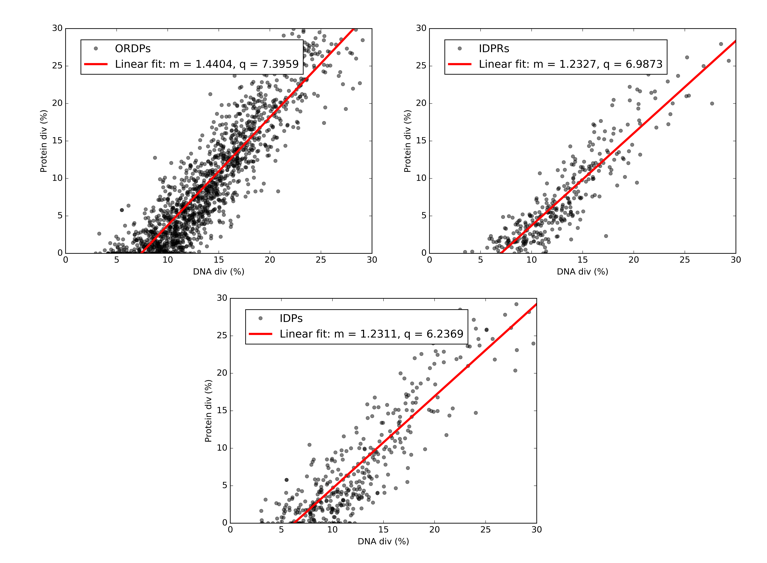

Canis familiaris	Tursiops truncatus

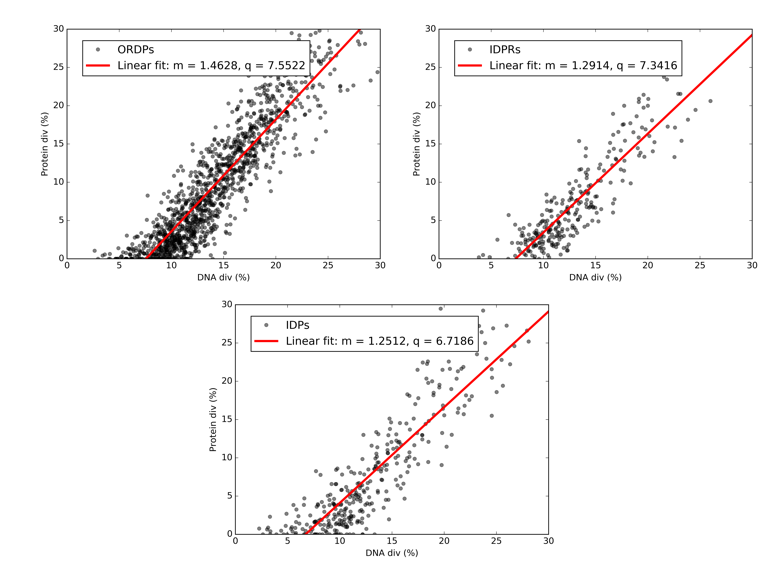

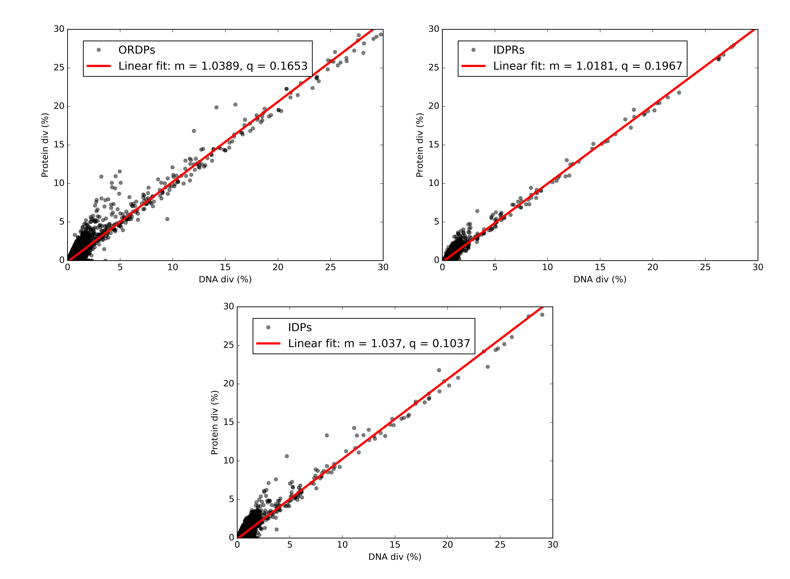

Bison bison bison	Mus spretus

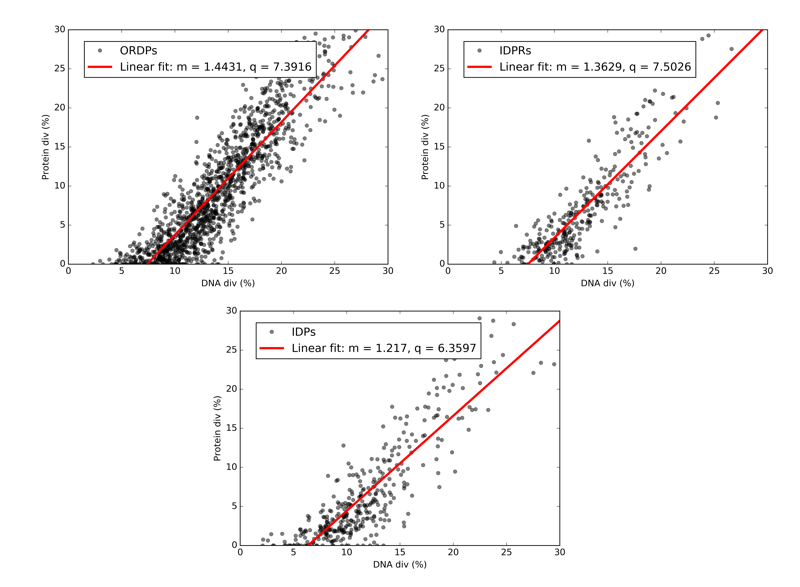

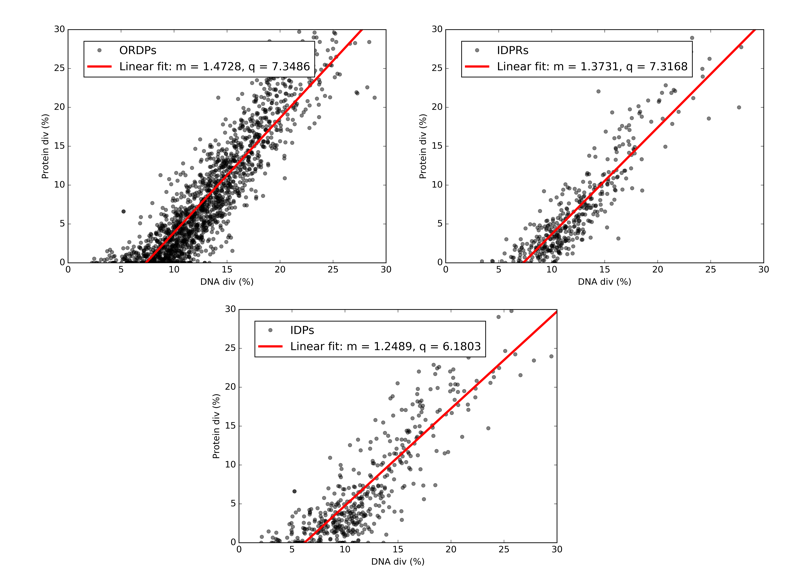

Sus scrofa	Felis catus

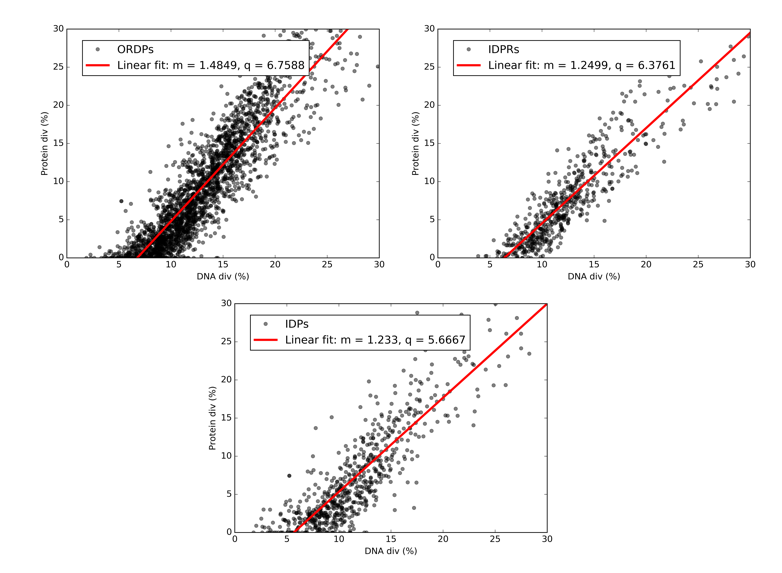

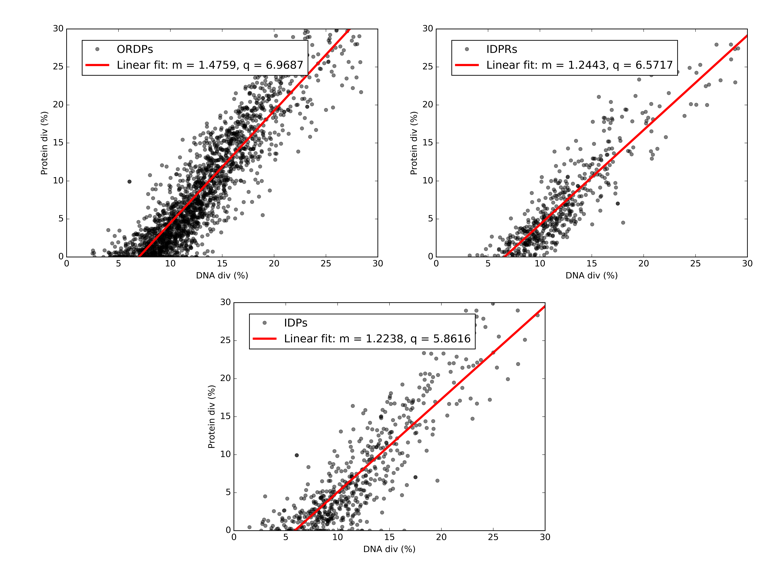

Castor canadensis	Macaca fascicularis

**Figure S2: DNA divergence Vs. Protein divergence plots.** Relationship between nucleotide (DNA div) and amino acid (Protein div) sequence divergence obtained by confronting progressively coding sequences of Mus Musculus (separated in ORDPs, IDPRs, and IDPs) with their homologs from 26 eukaryotes. Each point corresponds to an individual gene. In each panel, we report the best-fit line, together with the associated values of the slope (m) and the intercept (q) in the legend.
